# Long-term adaptation of prefrontal circuits in a mouse model of NMDAR hypofunction

**DOI:** 10.1101/2024.02.15.580421

**Authors:** Marion Ponserre, Tudor M Ionescu, Alessa A Franz, Serena Deiana, Niklas Schuelert, Thorsten Lamla, Rhîannan H Williams, Carsten T Wotjak, Scott Hobson, Julien Dine, Azar Omrani

## Abstract

Pharmacological approaches to induce N-methyl-D-aspartate receptor (NMDAR) hypofunction have been intensively used to understand the aetiology and pathophysiology of schizophrenia. Yet, the precise cellular and molecular mechanisms that relate to brain network dysfunction remain largely unknown. Here, we used a set of complementary approaches to assess the functional network abnormalities present in mice that underwent a 7-day subchronic phencyclidine (PCP 10mg/kg, subcutaneously, once daily) treatment. Our data revealed that pharmacological intervention with PCP affected cognitive performance and auditory evoked gamma oscillations in the prefrontal cortex (PFC) mimicking endophenotypes of some schizophrenia patients. We further assessed PFC cellular function and identified altered neuronal intrinsic membrane properties, reduced parvalbumin (PV) immunostaining and diminished inhibition onto L5 PFC pyramidal cells. A decrease in the strength of optogenetically-evoked glutamatergic current at the ventral hippocampus (HPC) to PFC synapse was also demonstrated, along with a weaker shunt of excitatory transmission by local PFC interneurons. On a macrocircuit level, functional ultrasound measurements indicated compromised functional connectivity within several brain regions particularly involving PFC and frontostriatal circuits. Herein, we reproduced a panel of schizophrenia endophenotypes induced by subchronic PCP application in mice. We further recapitulated electrophysiological signatures associated with schizophrenia and provided an anatomical reference to critical elements in the brain circuitry. Together, our findings contribute to a better understanding of the physiological underpinnings of deficits induced by subchronic NMDAR antagonist regimes and provide a test system for characterization of pharmacological compounds.

**Highlights:**

- Subchronic PCP treatment alters cognitive performance and evoked gamma synchronization in the prefrontal cortex.
- Subchronic PCP reduces number of parvalbumin-positive boutons and inhibitory synaptic transmission onto pyramidal neurons in layer 5/6 of the prelimbic cortex.
- Subchronic PCP reduces the strength of ventral hippocampal inputs to the medial prefrontal cortex.
- Frontostriatal circuits show robust dysconnectivity after subchronic PCP treatment. Abstract

## 1. Introduction

Schizophrenia features positive (e.g. hallucinations) and negative symptoms (e.g. social withdrawal, anhedonia), as well as cognitive impairments (e.g. attention, working memory deficits). Given the heterogeneity of the schizophrenia spectrum, developing animal models that accurately capture the intricate architecture of the disease has proven unrealistic. Nonetheless, animal models can faithfully reproduce some of the endophenotypes found in patients (Sigurdsson, 2016). The most studied endophenotypes include cognitive deficits (Arguello and Gogos, 2010); sensorimotor gating abnormalities (Powell et al., 2012); structural and functional dysregulation in the PFC and HPC as well as functional dysconnectivity between specific brain regions particularly within the frontostriatal networks (Meyer-Lindenberg and Weinberger, 2006; Tost et al., 2012); and dysregulation in neurochemical markers such as dopamine (Howes and Kapur, 2009), glutamate (Coyle, 2012; Javitt et al., 2012), and GABA (Lewis et al., 2012, 2005).

Within this framework, antagonists of the NMDAR have been at the forefront of preclinical and clinical research because they can recapitulate a full range of schizophrenia symptoms when given to healthy individuals (Anis et al., 1983; Krystal et al., 1994) and can model some of the aforementioned endophenotypes when administered to rodents (Frohlich and Horn, 2014; Lee and Zhou, 2019).

At the electrophysiological level, acute blockade of NMDAR elicits an increased activity of excitatory neurons and the release of glutamate in the medial PFC (mPFC) of mice and rats through a disinhibition mechanism (Gerhard et al., 2019; Homayoun and Moghaddam, 2007; Li et al., 2010; Moghaddam et al., 1997; Moghaddam and Adams, 1998; Suzuki et al., 2002). These changes resemble neurochemical disturbances present in patients in the acute phase of schizophrenia (Ben-Shachar et al., 2007; Callicott et al., 2000; Gur et al., 1995; Manoach et al., 2000, 1999; Soyka et al., 2005; Stone et al., 2009; Theberge et al., 2002) and gave rise to the glutamate hyperactivity theory (Moghaddam, 2003). This contrasts with the hypothesis of hypofrontality which has been consistently observed in schizophrenia patients (Hill et al., 2004; Townsend et al., 2022) and mimicked by long-term exposure to NMDA receptor antagonists (Cochran et al., 2003) in rodents. While a possible explanation for this “biphasic” response is that a hypoglutamatergic mPFC may arise after repetitive episodes of hyperglutamatergic states as seen after subchronic PCP exposure, the phenomenon is not fully understood, and the precise neuronal underpinnings are yet to be determined (Krzystanek and Pałasz, 2019). However, it is likely that the physiological disturbances seen in the PFC may require the effect of NMDAR blockade in other brain regions such as the HPC(Jodo, 2013; Nakazawa et al., 2017).

In rodents, the mPFC receives a dense monosynaptic glutamatergic projection from the ventral HPC (vHPC)(Condé et al., 1995; Marek et al., 2018; Parent et al., 2010). Importantly, decades of research in animal models and healthy individuals have upheld the concept that correlations in neural activity between these two structures play a crucial role in various cognitive functions (Sigurdsson and Duvarci, 2016). Meanwhile, it has become evident that reduced synchrony between them is a robust endophenotype of schizophrenia (Sigurdsson and Duvarci, 2016). Nevertheless, the relation of these deficits to the underlying anatomical and molecular impairments is still lacking.

In this study, we report a unique set of biomarkers that can define the subchronic PCP model from molecules to cells, and from local to macroscale circuits using state-of-the art technologies. We identify a pattern of disturbances centered around the mPFC and its local excitatory-inhibitory network. Our optogenetic-assisted circuit mapping data demonstrate a strong impairment of the vHPC input onto the mPFC. Finally, functional ultrasound highlights a robust dysconnectivity of frontostriatal circuits. All these features recapitulate the translational hallmark of NMDAR hypofunction. Our findings provide an anatomical circuit understanding of PFC dysfunction in the subchronic PCP model which is needed for improved characterization of new therapeutic concepts.

## 2. Materials and methods

### 2.1. Animals and drug treatment

Male C57BL/6JRj (Janvier Labs, later referred to as wild-type, WT, 5-7 weeks old) and C57BL/6-Tg(Pvalb-tdTomato)15Gfng (Taconic Biosciences A/S, 6-7 weeks old) mice were used for this study. All mice were group-housed (2-4) and kept in an animal facility under a 12-h light/dark cycle and all experiments were carried in the light phase except for the spontaneous alternation in T-Maze where mice were kept in a reverse light cycle and behavioral measurements were recorded during the dark phase. Food and water were provided ad libitum. All experimental procedures were designed to comply with the 3Rs (Tannenbaum and Bennett, 2015) and authorized by the Local Animal Care and Use Committee in compliance with local animal care guidelines, Association for Assessment and Accreditation of Laboratory Animal Care (AAALAC) regulations and the USDA Animal Welfare Act (19-025-G and 20-014-O). Different cohorts of mice were used for each experiment.

After habituation to housing condition, the mice (5-7 weeks of age) received either seven daily (Day 0 to Day 6) PCP (10mg/kg) or saline (Vehicle, Vh) injections subcutaneously (s.c). Vehicle and PCP-treated animals were housed separately (Fig. 1A).

**Fig. 1.**
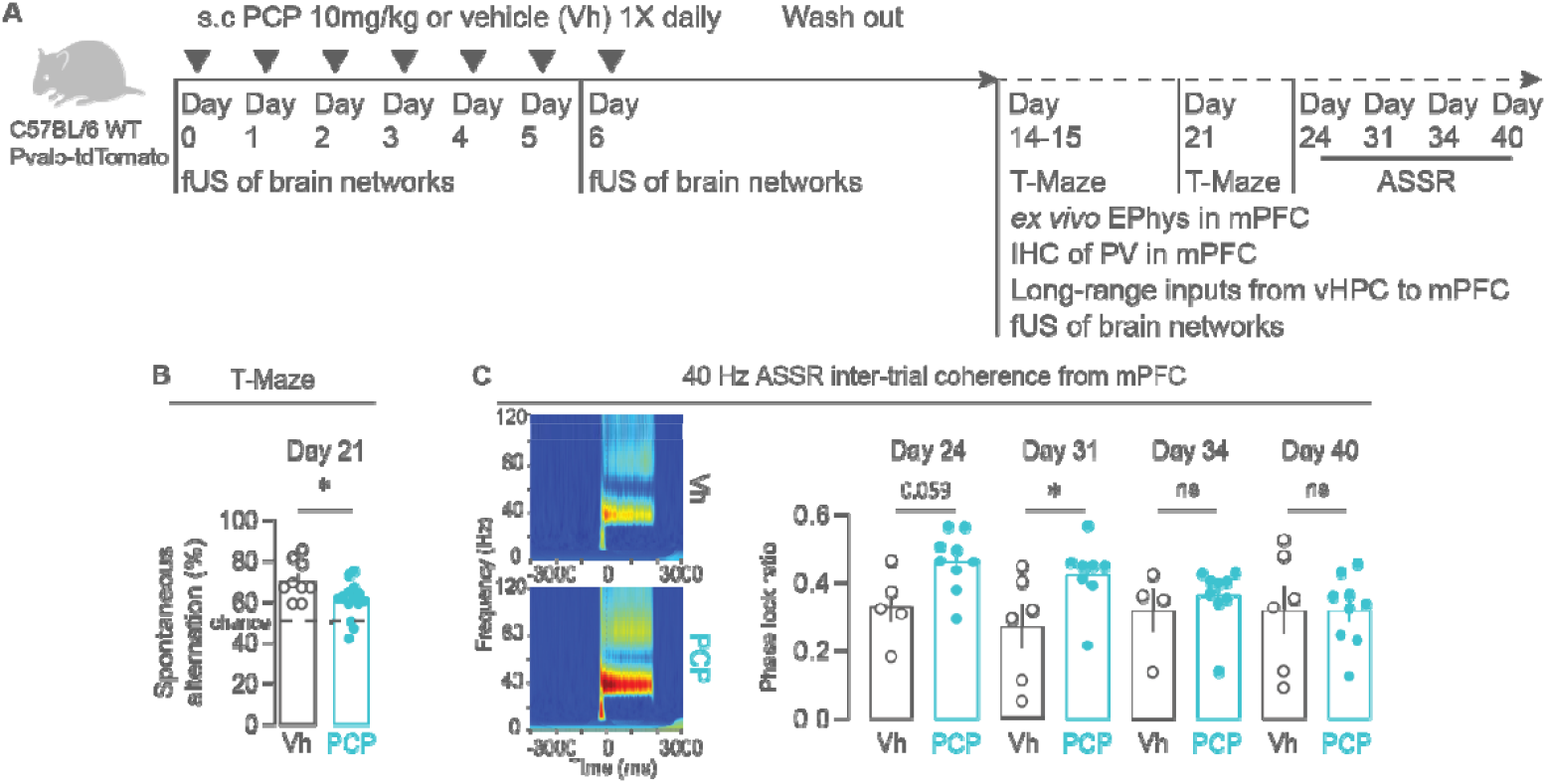
Mice subchronically treated with PCP display reduction in spontaneous alternation in the T-Maze and increase in ASSR phase-lock coherence in the mPFC. **(A)** Scheme showing the treatment regime used to study the effect of subchronic PCP on cognition using the T-Maze alternation task; on ASSR using electroencephalography (EEG); on local mPFC network using *ex vivo* electrophysiology (EPhys) and immunohistochemistry (IHC) of PV; on the long-range inputs from vHPC to mPFC using optogenetics; and on brain networks using functional ultrasound imaging (fUS). Different cohorts of mice were used for each experiment. **(B)** Percent of spontaneous alternation of vehicle- and PCP treated mice in the T-Maze task at day 21 (for Vh, mean spontaneous alternation: 70.3 ± 3.2 %, n= 9 mice; for PCP: 60.4 ± 8.9%, n= 15 mice; Mann-Whitney U test: U = 31.5, p = 0.03). (C) Heatmaps of phase-lock ratio (left) in 40 Hz ASSR from the mPFC of vehicle- and PCP treated animals and quantifications at day 24 (for Vh, mean ratio: 0.33 ± 0.046, n= 5 mice; for PCP: 0.462 ± 0.029, n= 9 mice; Mann-Whitney U test: U = 8, p = 0.0599) day 31 (for Vh, mean ratio: 0.27 ± 0.065, n= 6 mice; for PCP: 0.423 ± 0.035, n= 8 mice; Mann-Whitney U test: U = 7, p = 0.293); day 34 (for Vh, mean ratio: 0.32 ± 0.0627, n= 4 mice; for PCP: 0.3625 ± 0.030, n= 9 mice; Mann-Whitney U test: U = 12, p = 0.207) and day 40 (for Vh, mean ratio: 0.318 ± 0.071, n= 6 mice; for PCP: 0.318 ± 0.034, n= 9 mice; Mann-Whitney U test: U = 26, p = 0.964). Each dot is the quantification of a single animal.

### 2.2. Virus

The AAV8-hSyn-mChR2-mCherry was produced using a previously published protocol (Strobel et al., 2019) using high-density HEK293 cells seeded into 16-layer CELLdiscs and Calcium-phosphate based triple transfection of pHELPER, rep2/cap8 and pAAV-hSyn-mChR2-mCherry plasmids. The AAVs were purified by polyethylene glycol precipitation, iodixanol gradient centrifugation and ultrafiltration, as previously described (Strobel et al., 2015). Genomic titers were determined by qPCR using a hSyn promoter specific primer/probe set. AAV stocks were formulated in 1× PBS, 1 mM MgCl_2_, 2.5 mM KCl, 10% glycerol, 0.001% Pluronic F-68, pH 7.4. For experimental use, AAVs solution with a titer of 1.3×10 were diluted 1:1000 in PBS.

### 2.3. Stereotaxic surgeries

Male C57BL/6JRj mice were anaesthetized using isoflurane (induction, 3%; maintenance, 1.5%; CP-Pharma) in oxygen-enriched air and head fixed on a stereotaxic frame (model 1900, Kopf Instruments). Body temperature was maintained at 37°C using a heating pad. Eyes were protected with Bepanthen ointment, a local anaesthetic (bupivacaine 0.5% solution) was injected into the surgical site. After waiting 15 min, the surgical site was disinfected before the first incision.

For electrode implantation for ASSR recordings, five small holes were drilled into the skull. Coordinates for recording were -2.7 mm posterior and 4 mm lateral for the auditory cortex and 1.7mm posterior and 1mm lateral for the prefrontal cortex. The hole for the reference electrode was positioned above the cerebellum at +/-1.5 mm posterior to lambda. Gold-plated steel screws each with a connector cable and a metal pin were implanted into the cranium superficially of the dura mater. A socket with the connector pins to attach the Neurologger (TSE Systems) was implanted and fixed with dental cement.

For injection of AAV8-hSyn-mChR2-mCherry (see virus section above), 5 to 6 weeks old mice were bilaterally injected with 0.25 μl of virus (titer: 1.3×10^13^) in the vHPC by using the following coordinates calculated with respect to the bregma: -3.1 mm anteroposterior, 3.15 mm lateral, -4.6 to -4.5 mm ventral. Viral particles were delivered using glass pipettes (catalog #708707, Blaubrand intraMark) at a flow rate of 0.1 μl/min. After delivery of the virus, the pipette remained in the brain for 5 min to prevent spread of the virus. To alleviate postoperative pain, a long-acting analgesic (Meloxicam: 0.5 mg/kg subcutaneously) was given as a preventive measure, as well as on the first day after surgery and, depending on the animal’s condition, on the following three days. After the operation, the condition of the animals was monitored daily until they had recovered, and the wound had healed. Mice were treated with either PCP or vehicle (see Animals section above) 1 week after the stereotaxic surgery.

### 2.4. Spontaneous alternation in T-Maze

Spontaneous alternation in T-Maze was performed as previously described (Deiana et al., 2022) in male C57BL/6JRj mice (17-19gr at arrival), treated either with saline or PCP (see Animals and drug treatment section above). In brief, mice were tested in a single T-maze session, which consisted of 1 forced and a maximum of 14 free-choice trials or 10 minutes spontaneous exploration, whichever occurred first. Mice were tested 7 and 14 days post-PCP administration (day 14 and 21, as depicted in Fig. 1).

The percentage of spontaneous alternation over 14 free-choice trials was determined for each mouse and used as an index of working memory performance. Of note, this test is based on the foraging ethology of the mouse species; that is, mice have a natural tendency to explore novel places over known ones, as this behavior, often referred as shift-strategy, leads to a higher chance of finding food. This behavior requires intact working memory, as the animal must remember which arm it previously entered in order to alternate its choice.

### 2.5. ASSR recording

Detailed descriptions of the electrode implantation, setup specifications, recording procedure and analysis of the 40Hz ASSR protocol can be found in Schuelert et al. 2018 (Schuelert et al., 2018).

Male C57BL/6JRj mice were treated with either vehicle or PCP at the age of 38 to 41 days (see Animals and drug treatment section above) and implanted with cortical electrodes (age 50 to 53 days; see Stereotaxic surgery section above). Wireless recording of 40Hz auditory steady-state response (ASSR) was established using the Neurologger system (TSE Systems) and auditory stimuli generated with an audio generator (MED Associates Inc.). Mice were single-housed in sound-attenuated boxes (MED Associates Inc.) and presented with the auditory stimulus via a house speaker system. Using MED-specific software, the auditory stimulation protocols were programmed and a TTL signal was sent simultaneously with each tone to activate the LED infrared lights. An infrared sensor on the Neurologger recorded the infrared trigger signals and trigger signals were saved in the trigger channel. An ASSR session consisted of a periodic train of single white noise clicks at 40 Hz. Each train lasted 2 sec with an interval of 10 sec in between click trains. A total of 300 trains were presented. The intensity of the 40 Hz click train was adjusted to 85 +/-1.0 dB.

For each mouse, we recorded ASSR on day 24, day 31, day 34 and day 40 (Fig. 1). At the day of recording, Neurologgers were attached to the implanted connector pins 30 min before start of stimulation protocols. Vehicle and PCP-treated animals were always tested on the same day. Four recorded channels (bilateral auditory cortex electrodes, and bilateral prefrontal cortex electrodes) and one trigger channel were analyzed per animal.

The sampling rate of recording channels and trigger channels was set to 1000 Hz. The quality of EEG recordings was carefully checked and artifact segments removed. Data were analyzed using the Analyzer2 software package (BrainProducts GmbH, Munich, Germany).

For data analysis, data were segmented into 2.8 sec epochs with 400 msec pretrial period, 2 sec trial period and 400 msec post-trial period. A Morlet wavelet analysis was performed for the 40 Hz frequency layer to measure mean power and intertrial coherence (ITC) during the trial period. High ITC indicates a stable phase locking of brain responses to the auditory stimuli. ASSR power was expressed as the accumulated power of the 40 Hz layer during the stimulation phase of 2 sec. For the ITC wavelet, coefficients with complex numbers were computed and the phase-locking factor was determined by averaging the normalized phase synchronization across trials for every time point and frequency.

### 2.6. Immunohistology

Male C57BL/6JRj or Pvalb-tdTomato mice were treated with either vehicle or PCP (see Animals and drug treatment section above). On day 15, mice were deeply anesthetized by Pentobarbital (160 -200 mg/kg i.p), and transcardially perfused with 1x PBS (Gibco) followed by 4% paraformaldehyde (PFA) in 1x PBS (Electron Microscopy Sciences). Brains were post fixed in 4% PFA overnight at 4°C and afterwards transferred to 30% sucrose. Consecutive 30μm sections were prepared using a freezing stage sliding microtome (Epredia™ HM450). For free floating histology experiments, two sections per animal covering anterior and posterior PFC were blocked with 0.3% Triton X-100, 10% normal goat serum and 1% BSA (Dianova) in PBS for 2h at room temperature under steady agitation. Primary antibodies were diluted in 0.3% Triton X-100, 1% normal goat serum and 1% BSA (Dianova) (in the following referred to as carrier solution) and sections incubated overnight at 4°C under gentle agitation: rabbit anti-CaMKIIα (Millipore, #05-532, 1:200), guinea pig anti-Parvalbumin (Synaptic Systems, #195004, 1:500). Subsequently, sections were washed for about 2h with periodic exchange of washing solution in PBS containing 0.3% Triton X-100, 1% normal goat serum and 1%BSA (Sigma) and incubated with the following secondary antibodies diluted in carrier solution for 2h at room temperature and gentle agitation: goat anti-rabbit Alexa Fluor 488 (Invitrogen, #A11070, 1:1000), goat anti-guinea pig Alexa Fluor 647 (ThermoFisher, #A21450, 1:1000). After five washings steps of 15 min, sections were mounted in 24-well glass bottom plates (Sensoplate, Greiner), air dried and embedded with aqueous mounting medium containing DAPI (Fluoroshield™ with DAPI, Sigma). For the evaluation of protein levels in the prefrontal cortex, two coronal sections from the medial prefrontal cortex (Bregma 1.5 to 1.9)

### 2.7. High-throughput confocal microscopy and image analysis

Fluorescent images were obtained with an Opera Phenix (PerkinElmer) using the 63x objective in confocal mode. For each PFC section, 441 visuals fields (individual field size: 200 μm × 200 μm) were recorded, of which 20 fields were selected for the analysis of the posterior pre-limbic area, 28 fields for anterior pre-limbic and 32 fields for the analysis of the infra-limbic area according to the Allen Brain Atlas. Per individual field, a stack containing 5 planes separated by 1μm were imaged. Acapella Studio 5.5 software (PerkinElmer) was used to design a script for determining various read-out parameters. The DAPI staining was used to generate a nucleus mask for each field, which is necessary to identify cells positive for the respective fluorophores. PV-positive cells were determined by the fluorophore intensity in the nucleus and a global value for the background of the fluorescent signal. For PV positive cells, the nucleus/background ratio had to exceed a factor of 1.8. As the CaMKIIa soma staining was not consistently found within the nucleus region (as for PV cells), but rather in a peri-nuclear area, we adapted our detection algorithm for CaMKIIα -positive cells to identify cells as CaMKIIα -positive if the fluorescent intensity exceeds the factor 1.4 in the nucleus or within the peri-nuclear region. This region was defined as a 2.66 μm (14 pixel) ring outward the nucleus. Percental cell count was calculated as the number of positive cells per total DAPI cell count. Cell count and intensity were analyzed in maximum intensity projection mode. PV punctates onto CaMKIIα-positive cells as a readout for PV boutons were determined by the following synaptic parameters: the minimum spot intensity was set to 700 with a threshold factor for spot size of 900. PV spots were determined by a synapse intensity/background ratio factor of 1.3. PV spots on CaMKIIα-positive cells were determined by spots within the area of CaMKIIα-positive detected cells plus a ring of 2.28μm (12 pixel) around cells. Number of spots are normalized to the cell area (Nr. PV spots/μm^2^ ). Spots were analyzed per focal plane and data pooled.

### 2.8. *Ex vivo* electrophysiology

Male C57BL/6JRj or Pvalb-tdTomato mice were treated with either vehicle or PCP (see Animals and drug treatment section above). On day 14 (Fig. 1) mice were anesthetized (Pentobarbital 160 -200 mg/kg i.p), decapitated, and their brains were rapidly removed and transferred in ice-cold carbogenated (95% O2/5% CO2) cutting solution, containing (in mM), KCl 3, NaH_2_PO_4_ 1.25, Sucrose 218, Glucose 10, NaHCO_3_ 26, MgSO_4_ 10, CaCl_2_ 0.5 (pH 7.35). Coronal brain slices (250 μm) were prepared using a vibratome (Leica VT 1200⍰S). Slices were allowed to recover in oxygenated (95% O_2_ and 5% CO_2_) artificial cerebrospinal fluid (aCSF) containing the following chemicals (mM): NaCl 126, KCl 3, NaH_2_PO_4_ 1.25, Glucose 10, NaHCO_3_26, MgCl_2_ 1.3, CaCl_2_ 2.5. (pH 7.35) at 35⍰°C for at least 30⍰min, then kept at room temperature (22–23 °C) for at least another 30⍰min before experiments. Slices were transferred to the recording chamber and were continuously perfused at a flow rate of 2mL/min with oxygenated aCSF and maintained at room temperature. Slices were visualized using an upright epifluorescent microscope (Leica DM6 FS) with a 10× NA 0.22 or 25× NA 0.95 objective and equipped with infrared video microscopy and differential interference contrast (DIC) optics. For current clamp recording, a borosilicate glass pipette (3–5⍰MΩ) was filled with internal solution containing (in mM): K-gluconate 130, KCl 5, EGTA 0.5, HEPES 10, MgCl_2_ 2, Mg-ATP 4, Na-GTP 0.4, and K2-Phosphocreatine 10. The internal solution for voltage clamp recording contained (in mM): Cesium methanesulfonate (CsMes) 130, CsCl2 5, EGTA 0.6, HEPES 10, K2-Phosphocreatine 10, Mg-ATP 4 and Na-GTP 0.4. Signals were recorded at room temperature using an EPC10 patchclamp amplifier (HEKA). All signals were amplified and filtered at 2.9 KHz with a Bessel filter and digitized at 10 KHz. Access resistance was 6.5–25⍰MΩ and was monitored throughout the experiment. The membrane resistance and capacitance were monitored, although no criteria were set for these measures for inclusion. The slices of all virus-injected mice underwent post-hoc injection validation using immunohistochemistry.

For the analysis of spontaneous currents, pyramidal cells in L2/3 or L5/6 PL cortex were recorded in both C57BL/6JRj and Pvalb-tdTomato mice. Criteria to include cells were a leak current <100 pA at holding potential (−65 mV). Spontaneous inhibitory postsynaptic currents (sIPSCs) were recorded at 0 mV and spontaneous excitatory postsynaptic currents (sEPSC) were recorded at -65mV. sEPSC and sIPSC were recorded for 5 to 10 mins. Events were detected using Easy Electrophysiology (Easy Electrophysiology Ltd) and the template matching event detection algorithm using a correlation cut-off of 0.8.

For the analysis of evoked excitatory and inhibitory currents by ChR2-driven optogenetic stimulation, pyramidal cells in L5/6 PL cortex were recorded in virally injected C57BL/6JRj mice. Y3 filter cubes (Leica) was used to localized axons from vHPC neurons infected with the AAV8-ChR2 fused to mCherry. To stimulate the axons, 470-nm light pulses were applied with a CoolLed system (pE-300, CoolLED, UK) attached to the upright microscope (Leica DM6 FS). 1 ms pulses were used to evoke postsynaptic responses. Criteria to include cells in the analysis were an absolute leak current <150 pA at holding potential (−65 mV). We first recorded monosynaptic optogenetically evoked excitatory postsynaptic currents (oEPSC) at -65mV using a maximal light power (Max oEPSC amplitude). The power of the light was then reduced to reach an EPSC response with an amplitude corresponding to half of the amplitude reached with the max light power (Half power oEPSC amplitude). The same cell was then depolarized between -35mV and -40mV to record at the same time, the monosynaptic inward EPSC and the delayed di-synaptic evoked inhibitory postsynaptic current (oIPSC) using the same light power that produced half of the max oEPSC. Cells were excluded for further analysis if they showed a max oEPSC amplitude below 40 pA and/or if they did not show the di-synaptic oIPSC. An average of 5–10 traces were taken for analysis. Peaks were detected using the Curve Fitting function in Easy Electrophysiology.

For the analysis of intrinsic membrane properties, neurons of L5/6 PL cortex were recorded in both C57BL/6JRj and Pvalb-tdTomato mice. Neurons were accepted for recording if the resting membrane potential was more negative than −60 mV at break-in. For pyramidal cells recordings were included if action potentials evoked with an 800-ms step current injection displayed spike frequency accommodation (indicative of recording from pyramidal neurons, not interneurons). For PV interneurons, Y3 filter cubes (Leica) was used to localized tdTomato ^+^cells. Resting membrane potential was measured in bridge mode (I=0) immediately after obtaining whole-cell access and recordings were accepted if the cell was not spontaneously active at resting membrane potential (RMP) (the RMP was less than −55 mV). Input resistance was determined by injecting current steps of 800 ms with 20 pA increments starting from -100 to -60 pA; and calculated as the slope of the current-voltage plot using the Input Resistance function in Easy Electrophysiology. The voltage sag ratio was measured as a ratio of peak to steady-state voltage response to a hyperpolarizing current step of -300 pA for 800 ms and was calculated in Easy Electrophysiology using the Input Resistance function. The rheobase was estimated using a rising current ramp starting from -50 pA to 250 pA over 3.5s for pyramidal neurons and 5s for PV^+^ cells. To measure the firing frequency, firing was evoked by depolarizing current steps of 800 ms with 25pA increments starting from 0 to +225 pA. The properties of action potentials were recorded starting from the current injection step that triggered the first action potential. Properties were measured in Easy Electrophysiology for the first action potential only.

### 2.9. Functional ultrasound imaging

The experimental design and data preprocessing was identical to the one described in our previous work(Ionescu et al., 2023). For more details, please refer to the Supplementary Materials and Methods.

Male C57BL/6JRj were treated with either vehicle or PCP and measurements were done at day 0, day 6 and day 14 (Fig. 1). Functional connectivity (FC) matrices were generated for each animal by computing correlations between all pairs of the 12 preprocessed regional Power Doppler time courses. The generated Pearson’s r correlation coefficients were then transformed to Z-scores using Fisher’s Z-transformation and averaged to derive group-level connectivity matrices. Regional and connectivity strengths were computed by averaging the correlations of one region to all other regions. The first five minutes of all acquisitions were excluded to ensure the stability of the signals.

We first compared the FC of both readouts at baseline, between minutes 5 and 15 of the first measurement, briefly before applying the first PCP dose.

To compare the two cohorts at the day 0 timepoint and temporally characterize the effects of PCP on the PFC, we employed a sliding-window approach for dynamic FC analysis. We therefore computed FC in 10-minute windows slid by 30-second steps and extracted FC metrics for each window between the beginning and the end of each scan, and one matrix was generated for each animal and treatment group between 5 and 60 minutes after the start of the measurements. Using this approach, we identified the time-period for which the effect peaked, for which we present a comparison as a depiction of the maximum acute effect of PCP. To identify subchronic changes, we compared the FC of both cohorts for all edges, as well as the prefrontal FC strengths by analyzing the data acquired in the interval 5-60 minutes at days 7 and 14. For all comparisons, we tested for significance using two-sample t-tests between the two groups, which also yielded matrices of Z-scores, representing the magnitude of the significant differences between cohorts. We performed an FDR correction for multiple comparisons according to Benjamini-Hochberg (Benjamini and Hochberg, 1995). For dynamic FC at day 0, the correction was performed over time, all other corrections were performed over edges. To assess whether the spatial distribution of the effects remained stable between days 7 and 14, we further assessed the correlation of the generated Z-scores representing the statistical differences between the two cohorts at the two timepoints.

### 2.10. Statistical analysis

Analytical tests were performed with Prism (GraphPad) and Matlab for functional ultrasound. Outliers (if applicable) were determined by Grubb’s test (http://graphpad.com/quickcalcs/Grubbs1.cfm). Data distribution was tested for normality. When applicable, statistical tests were two-way ANOVAs or unpaired t-test. In case of not-normally distributed data, we used the Mann-Whitney test. Testing was always performed two-tailed with α = 0.05. Bar graphs show mean ± SEM, unless specified otherwise. ns = not significant, *p<0.05, ^**^p<0.01, ^***^p < 0.001, ^****^p < 0.0001 unless specified otherwise.

## 3. Results

### 3.1. Working memory and sensory processing deficits in a subchronic PCP model

We first addressed the effects of a subchronic PCP treatment on spontaneous alternation and cortical processing. C57BL/6JRj mice were injected once daily subcutaneously with PCP (10 mg/kg) or vehicle for 7 consecutive days (Fig. 1A). The animals were then subjected to a battery of in vivo and ex vivo tests either immediately after the first injection to capture the acute effect of the PCP treatment, after the last injection to see how these changes persisted and, or at several time points after the washout period (Fig. 1A).

To evaluate working memory performance, mice were tested in a T-maze spontaneous alternation task. In the first T-maze test (Day 14), PCP showed no effect (Supplemental Fig. 1A). On day 21 however, the performance of PCP-treated animals was significantly lower compared to vehicle-treated mice (Fig. 1B). To determine if this change was affiliated with altered cortical gamma power, we next assessed mPFC gamma oscillations via the auditory steady-state response (ASSR) paradigm. We measured the inter trial coherence (ITC) of evoked gamma which has been proposed as a biomarker of schizophrenia (Light et al., 2006; Roach et al., 2019; Sivarao et al., 2016), and found that subchronic PCP treatment significantly increased 40Hz ASSR ITC at day 31 (Fig. 1C) compared to vehicle-treated animals. We could also detect a trend toward increased 40Hz power (Supplemental Fig. 1A). The effect gradually diminishes over time, to reach the level observed in vehicle-treated animals by day 40 (Fig. 1C). Collectively, these findings indicate that sub-chronic PCP administration induces cognitive impairments and alters auditory evoked gamma oscillations in the PFC, recapitulating symptoms observed in schizophrenia patients.

### 3.2. Reduced parvalbumin expression in subchronic PCP model

EEG and behavioral experiments indicated that subchronic PCP treatment in mice may be modulating prefrontal circuitry. Since PV interneurons and NMDA receptors have been implicated both in generating gamma oscillations and in cognitive function (Gonzalez-Burgos et al., 2010; Gonzalez-Burgos and Lewis, 2012), we examined if PV interneurons in the PFC are affected by subchronic PCP treatment by assessing immunohistochemical markers at day 15 (Fig 1A).

Subchronic PCP did not affect the total number of identifiable prefrontal PV^+^cells within the mPFC (Fig. 2A-C and Supplemental Fig. 2A). Further examination revealed that immunofluorescent intensity of PV+ cells was affected, whereby the PV fluorescence intensity per cell was reduced when compared to vehicle-treated mice (Fig. 2D-E). This pattern was consistently repeated across the pre-limbic (PL) and infra-limbic (IL) cortical regions of the mPFC (Fig. 2A-D). Examination of PV-immunofluorescent boutons on CaMKIIα+ somata was also decreased, with a trend for reduced immunofluorescent intensity in the PL (Fig. 2F-H), but not in the IL (Supplemental Fig. 2D). Layer-specific anatomical evaluation revealed significantly reduced PV immunoreactivity in both layer 5/6 (L5/6, Fig. 2I) and layer 2/3 (L2/3, Supplemental Fig. 2E), yet the reduced PV+ bouton count and intensity was specific to L5/6 of PCP-treated animals (Fig. 2J and Supplemental Fig. 2F). In comparison, no change in either cell count or immunofluorescence intensity of CaMKIIα+ cells in PL after PCP treatment was observed (Supplemental Fig. 2A-C). Within the IL, there was a significant increase in CaMKIIα cells as a percentage of identifiable cells (Supplemental Fig. 2B). Overall, these data indicate that the prefrontal PV-driven inhibitory component is most prominently affected by subchronic PCP treatment, with the most pronounced reduction in PV immunoreactivity being observed in the deeper layer of the prelimbic area.

**Fig. 2.**
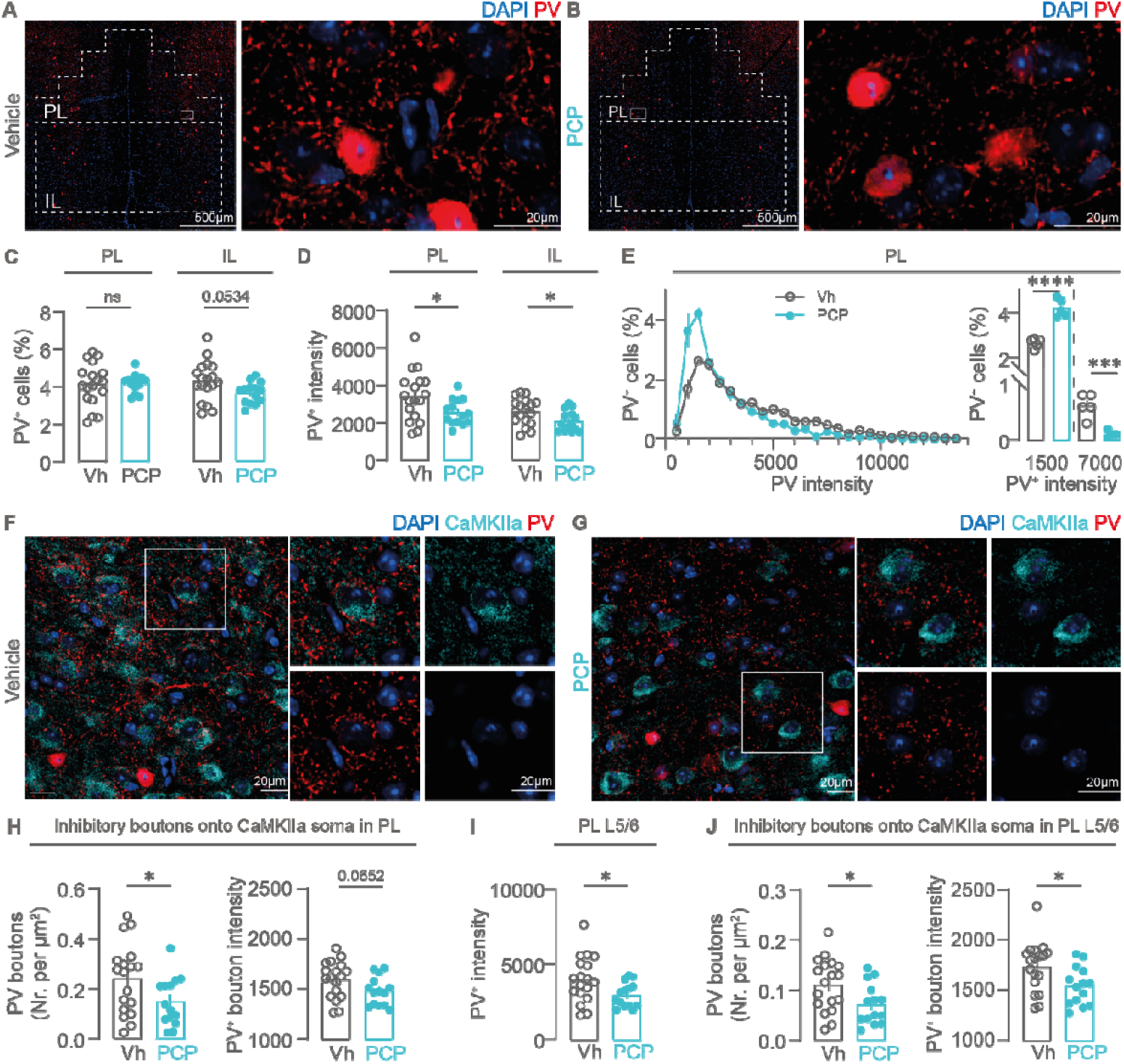
Subchronic PCP treatment reduces somatic and synaptic PV expression in the PL. **(A-B)** Representative confocal images of PV expression and DAPI positive cell nuclei in the mPFC area of vehicle-**(A)** and PCP-treated mice **(B)**. Dotted area indicate the imaged fields assigned to the PL and IL region of the PFC. **(C)** Percental count of PV ^+^cells in PL (left) and IL (right) (for Vh, mean total count: 4.148 ± 0.27, n=18 sections from 9 mice; For PCP, mean total count: 4.197 ± 0.13, n=14 sections from 7 mice; 2-tailed unpaired t-test: t (30) = 0.1529, p = 0.8795). **(D)** Somatic PV expression levels in PV^+^ cells in the PL of Vh- and PCP-treated mice (For Vh, mean PV intensity: 3433 ± 312.9, n= 18 sections from 9 mice; For PCP, mean PV intensity: 2576 ± 181, n=14 sections from 7 mice; 2-tailed t-test, t(30) = 2.196, p = 0.0359) and IL (For Vh, mean PV intensity: 2572 ± 165.8, n= 18 sections from 9 mice; For PCP, mean PV intensity: 2055 ± 146.2, n=14 sections from 7 mice; 2-tailed t-test: t(30) = 2.264 p = 0.0310). **(E)** Left: relative frequency distribution showing the PV intensity per PV cell (in nuclear cell region, see methods) from data quantified in (C). Right: Detail from left graph showing the percental amount of PV cells at low and high PV expression levels (Low PV levels: for Vh, mean PV %: 2.621 ± 0.09, n= 18 sections from 9 mice; For PCP, mean PV%: 4.188 ± 0.17, n=14 sections of 7 mice; 2-tailed unpaired t-test, t(8) = 7.831, p = < 0.0001. High PV levels: for Vh, mean PV %: 0.6048 ± 0.09, n= 18 sections from 9 mice; For PCP, mean PV%: 0.09025 ± 0.03, n=14 sections from 7 mice; 2-tailed unpaired t-test, t(8) = 5.301, p = 0.0007. Each data point shows mean of intensity bin, bin size = 100).**(F-G)** Representative confocal images of PV boutons on CaMKIIα pyramidal cells in PL of Vh-**(F)** and PCP-treated mice **(G)**. (H) Left: Density of inhibitory PV^+^ boutons on the soma of CaMKIIα excitatory cells in the PL in Vh- and PCP-treated mice (for Vh, mean density: 0.2401 ± 0.03, n=18 sections from 9 mice; For PCP, mean density: 0.1491 ± 0.03, n=14 sections from 7 mice; 2-tailed unpaired t-test: t(30) = 2.067, p = 0.0474). Right: PV fluorescence intensity of PV boutons on CaMKIIα^+^ somata in the PL (for Vh, mean intensity: 1593 ± 43.62, n=18 sections from 9 mice; For PCP, mean intensity: 1478 ± 38.23, n=14 sections from 7 mice; 2-tailed unpaired t-test: t(30) = 1.914, p = 0.0652). I PV immunoreactivity intensity in Layer 5/6 of PL (For Vh, mean PV intensity: 3999 ± 362.6, n=18 sections from 9 mice; For PCP, mean PV intensity: 2946 ± 219.7, n=14 sections from 7 mice; 2-tailed unpaired t-test: t (30) = 2.311, p = 0.0279) J Left: Density of inhibitory PV boutons on the soma of CaMKIIα excitatory cells in L5/6 PL of Vh- and PCP-treated mice (for Vh, mean density: 0.1113 ± 0.01, n=18 sections from 9 mice; For PCP, mean density: 0.072 ± 0.01, n=14 sections from 7 mice; 2-tailed unpaired t-test: t(30) =2.301, p = 0.0285). Right: PV fluorescence intensity of PV boutons on L5/6 CaMKIIα somata^+^ in the PL (for Vh, mean intensity: 1734 ± 58.86, n=18 sections from 9 mice; For PCP, mean intensity: 1539 ± 50.77, n=14 sections from 7 mice; 2-tailed unpaired t-test: t(30) = 2.426, p = 0.0215). Each data point represents mean measured per brain slice.

### 3.3. Compromised local mPFC network balance in subchronic PCP model

To explore whether repeated administration of PCP affects synaptic transmission in the mPFC, we recorded spontaneous inhibitory postsynaptic currents (sIPSCs) and excitatory postsynaptic currents (sEPSCs) from pyramidal cells of the mPFC from mice treated subchronic PCP (Fig. 1A). Given that PV bouton count and PV bouton intensity were specifically reduced in the PL of PCP-treated mice, synaptic currents were recorded in this area. We observed a strong and significant decrease in the frequency of sIPSCs in L5/L6 PL pyramidal neurons in PCP-treated compared with vehicle-injected mice (Fig. 3A and C), whereas sIPSC amplitudes were not affected (Fig. 3D). No differences were found in the frequency and amplitude of sEPSCs (Fig. 3C and D). In L2/3, sEPSC and sIPSCs remained unaffected (Fig. 3E-F) but the amplitude of sEPSCs was decreased (Fig. 3F). This suggests that the observed changes in inhibitory synaptic transmission may be more pronounced in the deeper layers of the prefrontal cortex.

**Fig. 3.**
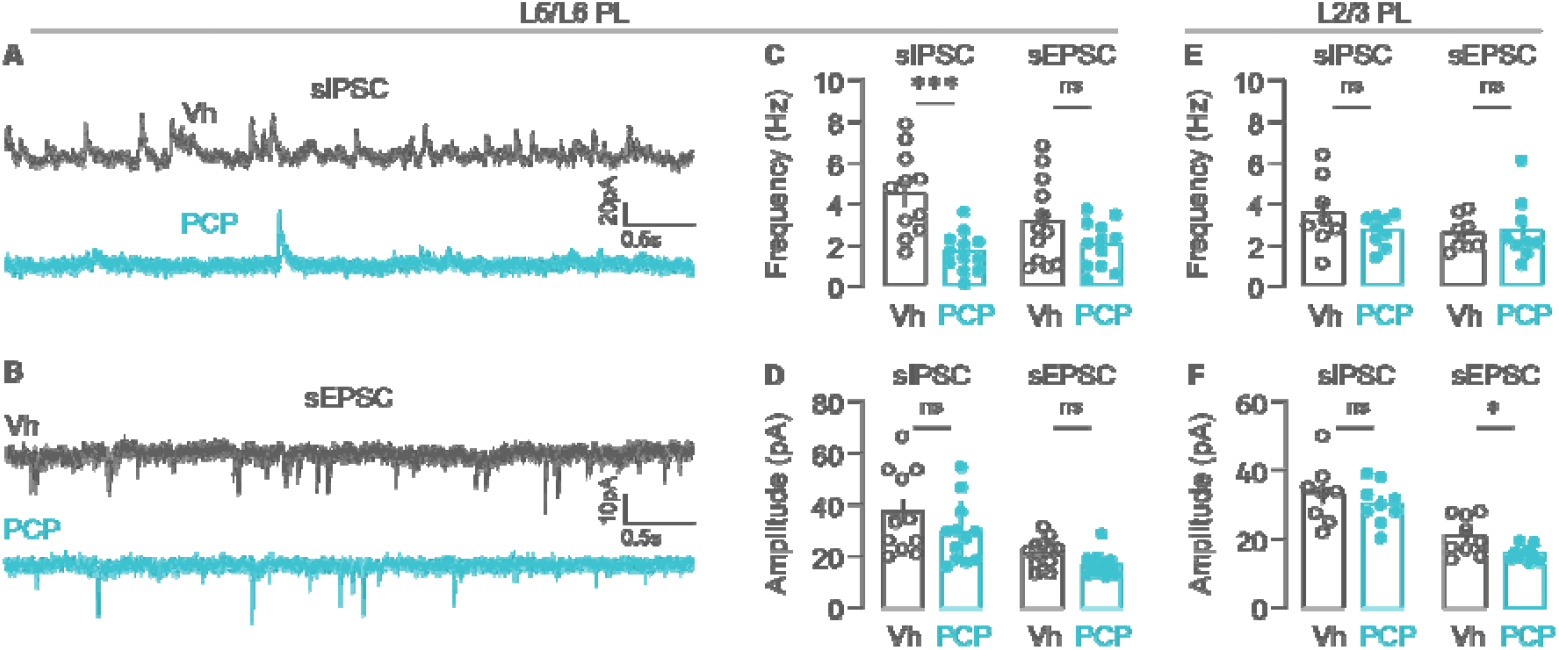
Inhibitory synaptic transmission onto pyramidal cells is decreased in PCP-treated animals. **(A-B)** Representative traces of sIPSCs **(A)** and sEPSC **(B)** from PC in L5/6 PL of Vh- and PCP-treated animals. **(C)** Mean frequency of sIPSCs and sEPSCs from PC in L5/6 PL of Vh- and PCP-treated animals (For sIPSCs, Vh mean frequency: 4.53 ± 0.61 Hz, n= 11 cells from 6 mice; PCP-treated mean frequency: 1.68 ± 0.28 Hz, n= 12 cells from 5 mice; 2-tailed unpaired t test: t(21) = 4.363, p = 0.0003; For sEPSCs, Vh mean frequency: 3.16 ± 0.6 Hz, n= 12 cells from 6 mice; PCP-treated mean frequency: 2.06 ± 0.33 Hz, n= 12 cells from 5 mice; 2-tailed unpaired t test: t(22) = 1.614, p = 0.1209). (D) Mean amplitude of sIPSCs and sEPSCs in PC in L5/6 PL of Vh- and PCP-treated animals (For sIPSCs, Vh mean amplitude: 37.2 ± 4.9 pA, n= 11 cells from 6 mice; PCP-treated mean amplitude: 29.3 ± 3.5 Hz, n= 12 cells from 5 mice; 2-tailed unpaired t test: t(21) = 1.344, p = 0.1932; For sEPSCs, Vh mean amplitude: 21.2 ± 1.7 pA, n= 12 cells from 6 mice; PCP-treated mean amplitude: 16.8 ± 1.34 pA, n= 12 cells from 5 mice; 2-tailed unpaired t test: t(22) = 2.036, p = 0.0539). (E) Mean frequency of sIPSCs and sEPSCs from PC in L2/3 PL of Vh- and PCP-treated animals (For sIPSCs, Vh mean frequency : 3.58 ± 0.6 Hz, n= 8 cells from 4 mice; PCP-treated mean frequency: 2.72 ± 0.25 Hz, n= 9 cells from 4 mice; 2-tailed unpaired t test: t(15) = 1.379, p = 0.1880; For sEPSCs, Vh mean frequency : 2.6 ± 0.29 Hz, n= 8 cells from 4 mice; PCP-treated mean frequency: 2.74 ± 0.51 Hz, n= 9 cells from 4 mice; 2-tailed unpaired t test: t(15) = 0.238, p = 0.8151). (F) Mean amplitude of sIPSCs and sEPSCs in PC in L2/3 PL of Vh- and PCP-treated animals (For sIPSCs, Vh mean amplitude : 33.28 ± 3.1 pA, n= 8 cells from 4 mice; PCP-treated mean frequency: 30.43 ± 2.0 pA, n= 9 cells from 4 mice; 2-tailed unpaired t test: t(15) = 0.7895, p = 0.4421; For sEPSCs, Vh mean amplitude : 21.1 ± 1.85 pA, n= 9 cells from 4 mice; PCP-treated mean amplitude: 15.9 ± 0.7 pA, n= 10 cells from 4 mice; 2-tailed unpaired t test: t(17) = 2.756, p = 0.014). Each dot is the quantification of a single cell.

We next examined the intrinsic membrane properties and excitability of both pyramidal neurons and PV interneurons in L5/6 of the PL. Surprisingly, cellular excitability of pyramidal neurons and PV interneurons in L5/6 of the PL were significantly lower (Supplemental Fig. 3A-C) and higher (Supplemental Fig. 3E-G) respectively, in PCP-treated mice. These changes were not associated with any change in passive and active membrane properties (Supplemental Table 1 and Table 2 and Supplemental Fig. 3D and H) although the resting membrane potential of PV neurons from PCP-treated mice was significantly more hyperpolarized (Supplemental Table 2). Taken together, these data reveal that subchronic PCP treatment results in changes to synaptic transmission, particularly impacting inhibitory synaptic inputs onto mPFC pyramidal cells.

### 3.4. Impaired hippocampal–PFC connectivity in a subchronic PCP model

Our data in the mPFC highlighted that PCP-induced differences were more pronounced in the deeper layers of the PL. In this region, there is abundant evidence that axons from long-range inputs present a laminar distribution with L5 being strongly innervated by vHPC efferents (Anastasiades and Carter, 2021). These findings prompted us to evaluate whether vHPC to mPFC inputs, which have been repeatedly involved in cognitive functions (Sigurdsson and Duvarci, 2016) are altered by PCP treatment. We used Channelrhodopsin-2(ChR2)–assisted circuit mapping (Petreanu et al., 2007) to characterize monosynaptic inputs from the vHPC to the mPFC in our subchronic PCP model. We transduced neurons in the CA1 of the vHPC with an AAV expressing ChR2-mCherry (Fig. 4A). Consistent with previous reports (Marek et al., 2018), we found that vHPC inputs innervated layers 2 to 5 of the infralimbic area (IL) and were segregated to L5 of the PL (Fig. 4A).

**Fig. 4.**
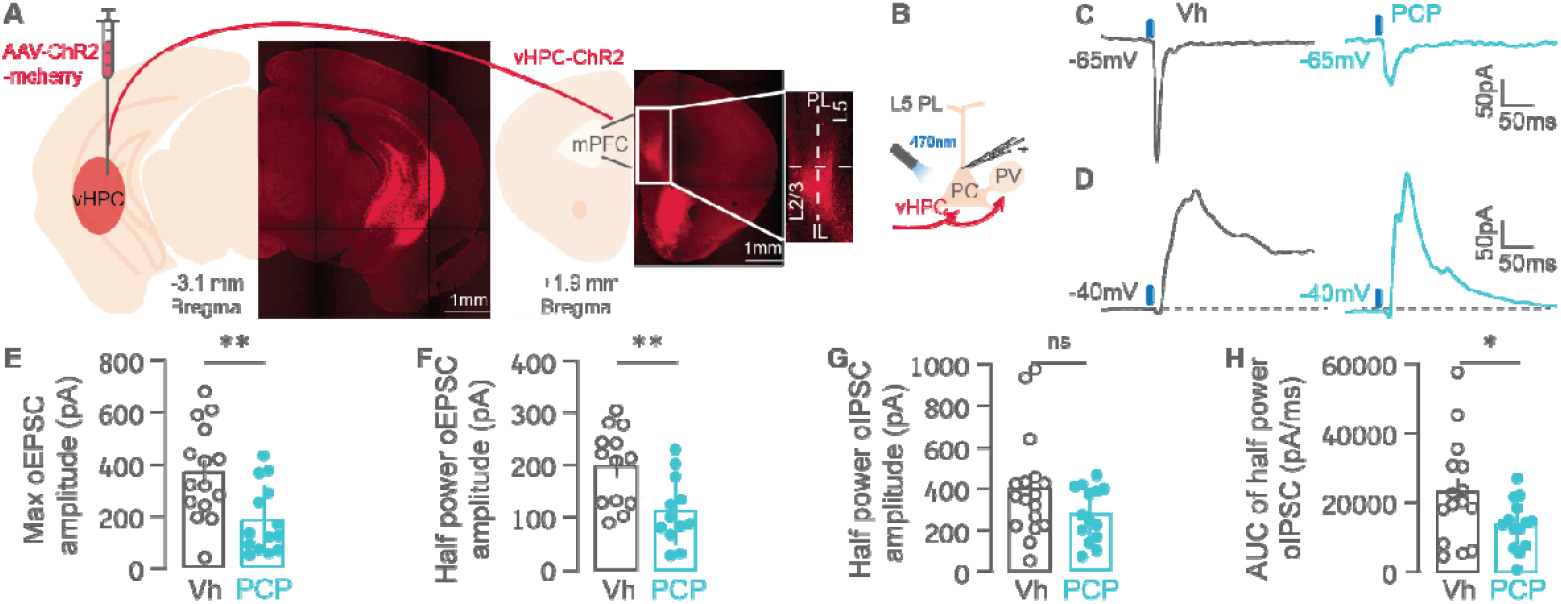
Monosynaptic excitatory transmission and feedforward inhibition are impaired between vHPC and mPFC in a subchronic PCP model. **(A)** Coronal images showing ChR2-mCherry expression at the injection site and labelling axon terminals in the mPFC. The inset (2x magnification) shows that vHPC inputs innervate L5 of the prelimbic and L2 to 5 of the infralimbic area. **(B)** Schematic of whole-cell recordings of pyramidal cells (PC) in L5 of PL. 470-nm light pulses were delivered to trigger action potentials in ChR2-expressing axon terminals to evoke postsynaptic responses in recorded cells. **(C-D)** Typical optogenetically-induced EPSC **(C)** oEPSC at holding potential of –65⍰mV) and IPSC **(D)** oIPSC at holding potential of –40⍰mV) in PC from vehicle- and PCP-treated animals. Synaptic responses in L5 PC in the PL show a monosynaptic EPSC followed by a feed-forward IPSC. **(E)** Max oEPSC amplitudes in PC of vehicle- and PCP-treated animals (For Vh, mean oEPSC max amplitude: 370.1 ± 46.4 pA, n= 15 cells from 10 mice; For PCP, mean oEPSC max amplitude: 187.2 ± 35.3 pA, n= 14 cells from 7 mice; 2-tailed unpaired t-test: t(27) = 3.1, p = 0.0044). **(F)** oEPSC amplitudes at a light power producing half of the max oEPSC in PC of vehicle- and PCP-treated animals (For Vh, mean half power oEPSC amplitude: 197.8 ± 20.5 pA, n= 13 cells from 10 mice; For PCP, mean half power oEPSC amplitude: 111.5 ± 64.6 pA, n= 12 cells from 7 mice; 2-tailed unpaired t-test: t(23) = 3.096, p = 0.0025). **(G)** oIPSC amplitudes at a light power producing half of the max oEPSC response in PC of vehicle- and PCP-treated animals (For Vh, mean oIPSC amplitude: 402.2 ± 60.5pA, n= 17 cells from 10 mice; For PCP, mean oIPSC amplitude: 279.3 ± 35 pA, n= 14 cells from 7 mice; 2-tailed unpaired t-test: t(29) = 1.661, p = 0.0537). **(H)** AUC of oIPSC at half max light power in PC of vehicle- and PCP-treated animals (For Vh, mean AUC of oIPSC: 23032 ± 3898 pA/ms, n= 15 cells from 10 mice; For PCP, mean AUC of oIPSC: 13648 ± 1903 pA/ms, n= 14 cells from 7 mice; 2-tailed unpaired t-test: t(27) = 2.115, p = 0.0219). Each dot is the quantification of a single cell.

Photostimulation of glutamatergic vHPC terminals (Fig. 4B) evoked excitatory postsynaptic currents (oEPSC) at a holding potential of -65 mV in pyramidal cells located in the L5 of PL (Fig. 4C). The amplitude of oEPSC were significantly smaller in PCP-treated animals (Fig. 4E-F). When pyramidal neurons were depolarized to – 40 mV, photostimulation of vHPC inputs produced a delayed disynaptic inhibitory postsynaptic current (oIPSC) that shunted the oEPSC (Fig. 4D). These findings align with an earlier study (Marek et al., 2018) indicating that pyramidal cells undergo feed-forward inhibition as a result of robust vHPC-driven activation of local fast-spiking PV interneurons (Fig. 4B). The vHPC-driven disynaptic IPSCs in pyramidal neurons were significantly smaller in PCP-treated mice as shown by reduced area under curve (AUC), although no significant change in the amplitude was observed (Fig. 4G-H).

Taken together our results show that monosynaptic excitatory inputs from vHPC onto pyramidal cells in L5 of the PL are largely reduced in PCP-treated animals. Furthermore, feed-forward inhibition onto pyramidal neurons is also decreased, likely as a result of a weaker monosynaptic excitatory input from the vHPC onto PV neurons in the mPFC and/or a deficit within the cortical microcircuit itself including decreased inhibitory drive from local PV neurons onto the pyramidal cells (see Fig. 4B).

### 3.5. Altered regional functional connectivity in a subchronic PCP model

Since subchronic PCP treatment induced deficits at cellular and microcircuit levels, we used fUS to probe if these modifications impact macroscale functional connectivity (Fig. 5). The probe was placed to record the signal of 12 ROIs, encompassing cortical and subcortical regions (Fig. 5) and we computed correlations between all pairs of the 12 regions as a readout for functional connectivity. The functional connectomes of both cohorts were not significantly different pre-injection (Fig. 5A and Supplemental Fig. 4A), the same applying for the prelimbic functional connectivity (FC) strength (Fig 5E). Acute PCP administration increased FC between several areas, most prominently involving the pallidum (Pal) and prelimbic cortex (Fig. 5B and Supplemental Fig. 4B), as well as enhanced global connectivity compared to vehicle (Fig. 5F and I). The increases in prelimbic FC started immediately after PCP administration, peaked between 25 and 35 minutes after application and were maintained until the end of the acquisition (Fig. 5I). In contrast, we saw robust decreases in frontal (including PL and anterior cingulate cortex (ACg)) and frontostriatal FC (including PL and caudate putamen area (CPu)) at both days 6 (Fig. 5C, G and Supplemental Fig. 4C) and days 15 (Fig. 5D, H and Supplemental Fig. 4D) following subchronic PCP treatment. While the subchronic changes at days 6 and 14 correlated significantly (Fig. 5J-K), they were more pronounced in the striatum, pallidum, and basolateral amygdala at day 14 (Fig. 5C-D).

**Fig. 5.**
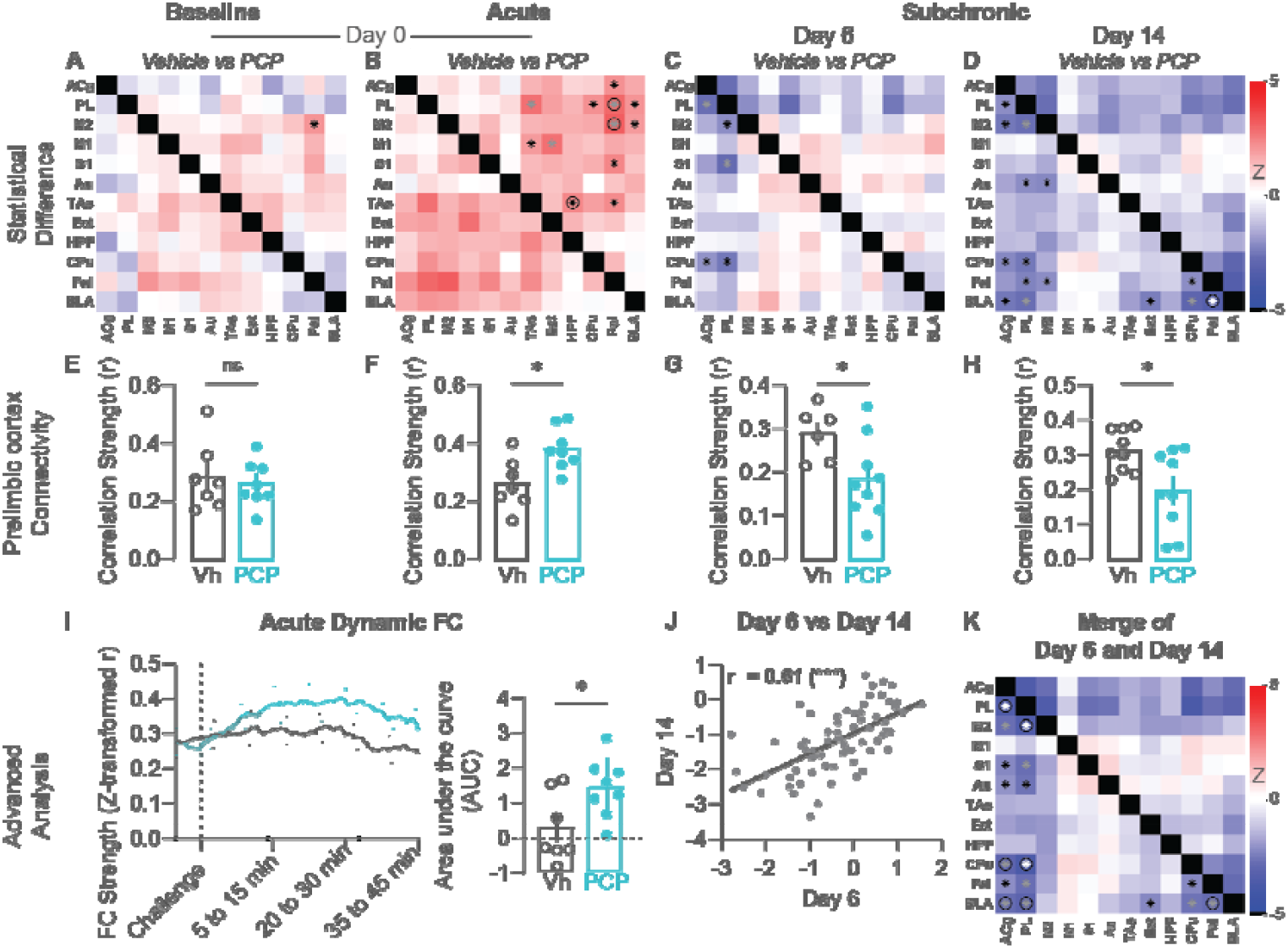
Functional connectivity analysis derived from fUS measurements upon acute and subchronic PCP treatment reveals abnormalities in brain networks. **(A-D)** Comparison of functional connectivity matrices of 12 regions covered by the acquired slice at **(A)** day 0 – baseline (minutes 5-15), **(B)** acute (minutes 25-35), **(C)** day 6 (minutes 5-60), and **(D)** day 14 (minutes 5-60). Matrices show the statistical differences (as Z-scores) between connectivity of the two cohorts. **(E-H)** Bar graphs at the bottom show connectivity strengths of the prelimbic cortex (averaged correlations of PL to all other regions) at day 0 (baseline **(E)** (For Vh, mean mPFC FC strength: 0.28 ± 0.12, n= 7 mice; For PCP, mean mPFC FC: 0.26 ± 0.07, n = 8 mice; 2-tailed unpaired t-test: t(13) = 0.48, p = 0.64) and **(F)** acute (For Vh, mean mPFC FC strength: 0.26 ± 0.09, n= 7 mice; For PCP, mean mPFC FC: 0.38 ± 0.07, n = 8 mice; 2-tailed unpaired t-test: t(13) = 2.94, p = 0.01)), **(G)** day 6 (For Vh, mean mPFC FC strength: 0.29 ± 0.06, n= 6 mice; For PCP, mean mPFC FC: 0.18 ± 0.09, n = 9 mice; 2-tailed unpaired t-test: t(13) = 2.41, p = 0.03), and (H) day 14 (For Vh, mean mPFC FC strength: 0.31 ± 0.05, n = 9 mice; For PCP, mean mPFC FC: 0.20 ± 0.11, n = 9 mice; 2-tailed unpaired t-test: t(16) = 2.69, p = 0.02). I Dynamic prelimbic FC analysis for the day 0 timepoint. A sliding window approach depicts the prelimbic FC over 10-minute blocks slid forward in 20-second steps between the beginning and the end of the scan for both cohorts. Right panel: corresponding areas under the curve for prelimbic connectivity strengths (For Vh, mean mPFC FC AUC: 0.28 ± 0.98, n = 7 mice; For PCP, mean mPFC AUC FC: 1.42 ± 0.85, n = 8 mice; 2-tailed unpaired t-test: t(13) = 2.39, p = 0.03). J Comparison of subchronic changes recorded at days 6 and 14 revealed strongly significant correlation between the respective Z-scores (r = 0.61, p < 0.0001). (K) Heatmap showing differences between both cohorts after merging the experiments performed at days 6 and 14. ^(*^black: p < 0.05, ^*^gray: p < 0.01, *white: p < 0.001, ^*^ in combination with O: p < 0.05 after FDR-correction, two-sample t-tests). Bar graphs show mean ± SEM, each dot is the quantification of a single animal. Graph in I show mean ± SD. Abbreviations: ACg = anterior cingulate, PL = prelimbic, M2 = secondary motor, M1 = primary motor, S1 = primary sensory, Au = auditory, TAs = temporal association, Ect = ectorhinal, HPF = hippocampal formation, CPu = caudate putamen, Pal = pallidum, BLA = basolateral amygdala.

Overall, using fUS we discovered that shifting from an acute to a subchronic NMDAR blockade prompted brain networks to transition from a hyperconnected to a hypoconnected state, with aberrances that were most pronounced within the prelimbic and frontostriatal circuits and which mimic the state observed in patients with schizophrenia(Anticevic et al., 2015b; Fornito et al., 2013).

## 4. Discussion

We propose the subchronic PCP model as a valuable tool for replicating the sensory processing impairment, specifically the alteration in the inter-trial coherence in a 40Hz ASSR paradigm (Powell et al., 2012), as well as the cognitive dysfunction (Arguello and Gogos, 2010) seen in schizophrenia. Moreover, this treatment regime mirrored the reduced PV expression and local network alterations in the mPFC (Lewis et al., 2012, 2005) associated with the disorder. Finally, it reproduced functional dysconnectivity between the hippocampus and prefrontal cortex, as well as within the frontostriatal networks (Meyer-Lindenberg and Weinberger, 2006; Tost et al., 2012).

Deficits in working memory are a dominant cognitive symptom in schizophrenia and it has been shown that PFC-dependent working memory depends on intact transmission from PV interneurons (Murray et al., 2015). In line with these studies, we found that PCP-treated animals performed significantly worse in the T-maze spontaneous alternation task at day 21. As an animal must recall which arm it visited in the previous trial, working memory must be intact to elicit normal spontaneous alternation. Hence, the observed impairment in spontaneous alternation following PCP exposure aligns with past studies, confirming that chronic antagonism of NMDAR adversely impacts performance related to working memory (d’Isa et al., 2021; Driesen et al., 2013; Lisman et al., 1998; Rezvani, 2006).

Besides the noted cognitive dysfunction, we also identified aberrances in the synchronization of the PFC network in the 40 Hz gamma frequency band in response to steady-state auditory stimulation. This was evidenced by an increase in ASSR phase-lock ratio after subchronic PCP treatment which resolved over time. These findings align with a previous study (Leishman et al., 2015) conducted in rats and showing that both acute and subchronic administration of PCP can lead to an increase in the synchrony and power of 40Hz ASSR. Schizophrenia patients exhibit deficits in the ability to generate gamma frequency oscillations in response to ASSR, making the 40Hz ASSR a biomarker that can be measured in both preclinical (Sivarao et al., 2016) and clinical settings (Light et al., 2006; Roach et al., 2019). However, EEG studies of ASSR in schizophrenia patients have reported conflicting findings, with some showing a reduction (Reilly et al., 2018), and others an increase (Hamm et al., 2012; Kim et al., 2019) in ASSR; a discrepancy which may be attributed to variations in stimulus parameters(Kim et al., 2019). Importantly, by optogenetically inhibiting PV-interneurons, 40Hz ASSR ITC could be reduced (Toader et al., 2020), confirming the relevance of functional PV interneurons for 40 Hz ASSR (Balla et al., 2020; Gonzalez-Burgos et al., 2010). In our model, it is possible that the increase in excitability of PV ^+^cells as seen using *ex vivo* recordings of these neurons could account for the *in vivo* increase in ASSR phase-lock ratio. In the basal forebrain for instance, phase-locked excitation of PV neurons^+^ in advance of 40 Hz auditory stimuli have been shown to enhance the power of cortical ASSR responses (Hwang et al., 2019).

We further examined PV protein expression using immunohistochemistry and discovered that subchronic PCP administration decreased PV protein expression in the mPFC. In humans, lower levels of PV mRNA expression have been consistently observed in the cortex and hippocampus of schizophrenia patients (Benes et al., 1991; Fung et al., 2010; Hashimoto et al., 2003; Mellios et al., 2009; Zhang and Reynolds, 2002). Similar findings have been reported in rodents following acute and subchronic treatments with NMDAR antagonists (Behrens et al., 2007; Keilhoff et al., 2004; Romón et al., 2011; Zhou et al., 2015). More broadly, a converging body of literature has stressed the vulnerability of PV^+^ cells in various brain disorders (Marín, 2012). In our subchronic PCP model, lower PV expression was associated with a reduced number of PV boutons onto pyramidal neurons. This supported our *ex vivo* electrophysiology study showing decreased sIPSCs frequency onto L5 pyramidal cells. Surprisingly however, PV neurons were more excitable in PCP-treated mice which may suggest a compensatory mechanism to overcome the decreased inhibitory tonus onto pyramidal cells. It is for instance possible that the observed decrease in PV protein expression contributes to increase the excitability of PV neurons. In line with this, initial research has revealed that complete absence of PV enhances the excitability of fast spiking interneurons (Orduz et al., 2013). On the other hand, more recent work has described that partial PV downregulation did not change the intrinsic properties of fast spiking PV^+^cells but was sufficient to reduce local inhibitory transmission onto pyramidal neurons (Caballero et al., 2020). Further research is required to clarify the relationship between increased PV ^+^cells excitability and decrease inhibitory inputs onto pyramidal neurons. Importantly, the similarity of changes in PV expression in schizophrenia and after PCP subchronic administration adds construct validity to our model. Furthermore, the alterations in both PV interneurons and pyramidal cells functionality likely results in a reduction of the coordinated activity of large brain networks, a core endophenotype of schizophrenia.

Reduced synchrony between the HPC and PFC has been repeatedly reported in animal models using *in vivo* local field potential recordings. This dysfunction is evident across various behavioral conditions and has been identified using diverse analytical methods (Sigurdsson and Duvarci, 2016). Here, we aimed to examine how abnormal ventral hippocampal-prefrontal coordination in a subchronic PCP model manifested itself at the level of synaptic transmission between these two regions. We used optogenetic technology to specifically target vHPC inputs projecting onto mPFC pyramidal cells. In line with previous studies (Marek et al., 2018), activation of vHPC inputs in the mPFC induced excitation followed by a delayed feed-forward inhibition of the pyramidal cells. In PCP-treated animals, both excitatory and inhibitory responses were reduced. While the net effect on the PFC network of these disruptions is yet to be tested, our results align with previous observations of reduced influence of the HPC on the PFC in schizophrenia (Benetti et al., 2009). In addition, given that prefrontal neurons are phase-locked to hippocampal theta oscillations (Buzsáki, 2002; Sigurdsson and Duvarci, 2016), it is expected that synchrony between the two regions during various cognitive tasks will be affected. This hypothesis is supported by previous studies using genetic mouse models showing impairments in hippocampal-prefrontal communication that were associated with diminished branching (Mukai et al., 2015) and reduced probability of neurotransmitter release at the hippocampal–mPFC glutamatergic synapse (Dawson et al., 2015). Nonetheless, further experiments are required in our subchronic PCP model to understand how the identified alterations at the vHPC to mPFC synapses will influence the firing rate of prefrontal pyramidal cells.

In order to detect changes in brain network communication beyond HPC-mPFC circuit, we used functional ultrasound imaging. Our fUS data highlighted a transition from hyperconnected to hypoconnected PFC networks in PCP-treated animals. In accordance with our earlier study using ketamine (Ionescu et al., 2023), we observed strong, global FC and cerebral blood volume (CBV) surges after acute NDMAR antagonist treatment. These effects are also in line with acute effects of NMDAR antagonists on FC reported using ketamine in humans (Anticevic et al., 2015a; Kotoula et al., 2021) and on CBV in mice (Masaki et al., 2019). Our data also confirms earlier work performed with BOLD-fMRI showing that PCP, when given acutely in mice, induces global hyperconnectivity within the first 20 minutes after injection (Montani et al., 2021). The global increases in FC have been hypothesized to be driven by a state of excessive glutamatergic signaling (Anticevic et al., 2015a). Specifically for the PFC, the hyperconnectivity induced by acute NMDAR antagonists has been proposed to partly mimic early-course schizophrenia (Anticevic et al., 2015a). Interestingly, the same study reported that the PFC exhibited global hypoconnectivity in chronic schizophrenia patients (Anticevic et al., 2015a) which was also a robust feature of the functional connectome after subchronic PCP treatment here. While our ex vivo electrophysiological recordings were conducted at time points corresponding to an hypoconnected state as shown using fUS, it would be intriguing to elucidate the precise neuronal mechanisms that govern the transition from hyperconnected to hypoconnected networks, especially between the prelimbic cortex and its long-range inputs.

Frontostriatal hypoconnectivity has also been consistently reported as a major hallmark of functional (Dandash et al., 2014; Karcher et al., 2019; Orliac et al., 2013) and structural (Ellison-Wright and Bullmore, 2009) connectomic disruption in schizophrenia. Reductions in frontostriatal connectivity have been shown in first-degree relatives of schizophrenic patients (Fornito et al., 2013) and these reductions have been observed to return to normal levels in patients exhibiting better outcomes following medication (Sarpal et al., 2015). On neurotransmitter level, the observed frontostriatal dysconnectivity may be at the crossroads of both glutamatergic and dopaminergic malfunction in the two regions, prefrontal hypoactivity being able to lead to a hyperdopaminergic state in the striatum (Meyer-Lindenberg et al., 2002). A recent preclinical study on the effect of an NMDAR antagonist on dopamine synthesis showed that subchronic ketamine treatment strongly increased striatal dopamine synthesis and locomotor activity (Kokkinou et al., 2021). Interestingly, both of these phenotypes could be rescued by increasing prefrontal PV interneuron activity.

The reduction in anterior cingulate FC we observed here may additionally mimic disturbed integrities of resting-state networks. The anterior cingulate cortex anchors both the salience network and, together with the prefrontal cortex, forms the anterior default mode network (DMN) in rodents. Disruptions in both networks have been consistently found in literature in schizophrenia patients (Karcher et al., 2019; Orliac et al., 2013).

A major limitation of our study is the lack of whole-brain coverage of our fUS acquisitions. For the prefrontal cortex, it has been shown that FC of neighboring areas can display different connectomic changes in schizophrenia (Chechko et al., 2018). Therefore, since we only acquired one slice of the PFC, it is difficult to determine whether a slightly different positioning of the probe could have revealed a different pattern of alterations, perhaps also better capturing a fronto-hippocampal dysconnectivity. Additionally, the use of sedated animals may affect the observed readouts in an unpredictable manner (Jonckers et al., 2015, 2014). It is plausible that fronto-hippocampal dysconnectivity may be better captured during a cognitive task condition, than during resting-state ^87^.

## 5. Conclusions

In conclusion, our functional ultrasound imaging, which measured functional connectivity following both acute and subchronic PCP injections, along with previous literature using the acute and subchronic NMDAR antagonist models, suggest a dynamic reorganization of excitatory and inhibitory inputs at various time points. This adds complexity to interpreting our findings. Despite this, our data offers valuable insights into these time-dependent observations. Notably, upon acute PCP injection, we detected a hyperconnected PFC network which transitioned to an hypoconnected state after subchronic treatment. This was concurrent to disruptions in the local PFC network which manifested as changes in neuronal intrinsic membrane properties, and reduced inhibition of pyramidal cells. We additionally uncovered a decrease in the strength of evoked glutamatergic current between the HPC and the PFC. Future research should explore how the dynamic changes, from acute to subchronic PCP treatment, may mirror the neurobiological alterations associated with the progression of schizophrenia. The potential of pharmacological intervention to reverse these abnormalities in the prodromal phase and prevent further progression offers a promising avenue.

## Abbreviations

ACg: Anterior cingulate
ASS: RAuditory steady-state response
AuA: uditory
AUC: Area under the curve
BLA: Basolateral amygdala
ChR2: Channelrhodopsin 2
CPu: Caudate putamen
DMN: Default mode network
Ect: Ectorhinal
EEG: Electroencephalography
FC: Functional connectivity
fUS: Functional ultrasound imaging
HPC: Hippocampus
HPF: Hippocampal formation
IHC: Immunohistochemistry
IL: Infralimbic
ITC: Intertrial coherence
M1: Primary motor
M2: Secondary motor
mPFC: Medial PFC
NMDAR: N-methyl-D-aspartate receptor
oEPSC: Optogenetically evoked excitatory postsynaptic current
oIPSC: Optogenetically evoked inhibitory postsynaptic current
Pal: Pallidum
PCP: Phencyclidine
PFC: Prefrontal cortex
PL: Prelimbic
PV: Parvalbumin
RMP: Resting membrane potential
S1: Primary sensory
sEPSC: Spontaneous excitatory postsynaptic currents
sIPSC: Spontaneous inhibitory postsynaptic currents
Tas: Temporal association
Vh: Vehicle
vHPC: Ventral hippocampus

## CRediT authorship contribution statement

AO and MP conceived the study. MP designed, performed, and analyzed data from ex vivo electrophysiology experiments and stereotaxic surgeries; TI designed, performed, and analyzed data from fUS imaging experiments; AF designed, performed, and analyzed data from immunohistology experiments; SD designed, and analyzed data from T-Maze experiments; NS designed, and analyzed data from ASSR recordings, TL produced the AAV-hSyn-mChR2-mCherry virus; RHW, CTW, SH, and JD provided critical insight. MP, TI and AF drafted the manuscript with inputs from all other co-authors. AO supervised the project.

## Funding

This work was supported by Boehringer Ingelheim Pharma & Co. KG.

## Declaration of competing interest

All authors are employees of Boehringer Ingelheim Pharma & Co. KG.

## Acknowledgments

The authors would like to thank Dr. Amarender Bogadhi for help with data analysis in MATLAB; Dr. Bernd Igl for statistical consulting; Johannes Freudenreich, Dr. Stefan Jäger, Nancy Kötteritzsch, Matthias Heil, Niklas Lindner, Aileen Reich, and Carmen Weiss for their excellent technical contributions; and Dr. Benjamin Strobel for providing the the AAV8-hSyn-mChR2-mCherry.

## Appendix A

Supplementary data

## References

Anastasiades, P.G., Carter, A.G., 2021. Circuit organization of the rodent medial prefrontal cortex. Trends Neurosci 44, 550–563. 10.1016/j.tins.2021.03.006

Anis, N.A., Berry, S.C., Burton, N.R., Lodge, D., 1983. The dissociative anaesthetics, ketamine and phencyclidine, selectively reduce excitation of central mammalian neurones by N□methyl □aspartate. Br. J. Pharmacol. 79, 565–575. 10.1111/j.1476-5381.1983.tb11031.x

Anticevic, A., Corlett, P.R., Cole, M.W., Savic, A., Gancsos, M., Tang, Y., Repovs, G., Murray, J.D., Driesen, N.R., Morgan, P.T., Xu, K., Wang, F., Krystal, J.H., 2015a. N-Methyl-D-Aspartate Receptor Antagonist Effects on Prefrontal Cortical Connectivity Better Model Early Than Chronic Schizophrenia. Biol. Psychiatry 77, 569–580. 10.1016/j.biopsych.2014.07.022

Anticevic, A., Hu, X., Xiao, Y., Hu, J., Li, F., Bi, F., Cole, M.W., Savic, A., Yang, G.J., Repovs, G., Murray, J.D., Wang, X.-J., Huang, X., Lui, S., Krystal, J.H., Gong, Q., 2015b. Early-Course Unmedicated Schizophrenia Patients Exhibit Elevated Prefrontal Connectivity Associated with Longitudinal Change. J. Neurosci. 35, 267–286. 10.1523/jneurosci.2310-14.2015

Arguello, P.A., Gogos, J.A., 2010. Cognition in Mouse Models of Schizophrenia Susceptibility Genes. Schizophr. Bull. 36, 289–300. 10.1093/schbul/sbp153

Balla, A., Ginsberg, S.D., Abbas, A.I., Sershen, H., Javitt, D.C., 2020. Translational neurophysiological biomarkers of N-methyl-d-aspartate receptor dysfunction in serine racemase knockout mice. Biomark. Neuropsychiatry 2, 100019. 10.1016/j.bionps.2020.100019

Behrens, M.M., Ali, S.S., Dao, D.N., Lucero, J., Shekhtman, G., Quick, K.L., Dugan, L.L., 2007. Ketamine-Induced Loss of Phenotype of Fast-Spiking Interneurons Is Mediated by NADPH-Oxidase. Science 318, 1645–1647. 10.1126/science.1148045

Benes, F.M., McSparren, J., Bird, E.D., SanGiovanni, J.P., Vincent, S.L., 1991. Deficits in Small Interneurons in Prefrontal and Cingulate Cortices of Schizophrenic and Schizoaffective Patients. Arch. Gen. Psychiatry 48, 996–1001. 10.1001/archpsyc.1991.01810350036005

Benetti, S., Mechelli, A., Picchioni, M., Broome, M., Williams, S., McGuire, P., 2009. Functional integration between the posterior hippocampus and prefrontal cortex is impaired in both first episode schizophrenia and the at risk mental state. Brain 132, 2426–2436. 10.1093/brain/awp098

Benjamini, Y., Hochberg, Y., 1995. Controlling the False Discovery Rate: A Practical and Powerful Approach to Multiple Testing. J Royal Statistical Soc Ser B Methodol 57, 289– 300. 10.1111/j.2517-6161.1995.tb02031.x

Ben-Shachar, D., Bonne, O., Chisin, R., Klein, E., Lester, H., Aharon-Peretz, J., Yona, I., Freedman, N., 2007. Cerebral glucose utilization and platelet mitochondrial complex I activity in schizophrenia: A FDG-PET study. Prog. Neuro-Psychopharmacol. Biol. Psychiatry 31, 807–813. 10.1016/j.pnpbp.2006.12.025

Buzsáki, G., 2002. Theta Oscillations in the Hippocampus. Neuron 33, 325–340. 10.1016/s0896-6273(02)00586-x

Caballero, A., Flores-Barrera, E., Thomases, D.R., Tseng, K.Y., 2020. Downregulation of parvalbumin expression in the prefrontal cortex during adolescence causes enduring prefrontal disinhibition in adulthood. Neuropsychopharmacology 45, 1527–1535. 10.1038/s41386-020-0709-9

Callicott, J.H., Bertolino, A., Mattay, V.S., Langheim, F.J.P., Duyn, J., Coppola, R., Goldberg, T.E., Weinberger, D.R., 2000. Physiological Dysfunction of the Dorsolateral Prefrontal Cortex in Schizophrenia Revisited. Cereb. Cortex 10, 1078–1092. 10.1093/cercor/10.11.1078

Chechko, N., Cieslik, E.C., Müller, V.I., Nickl-Jockschat, T., Derntl, B., Kogler, L., Aleman, A., Jardri, R., Sommer, I.E., Gruber, O., Eickhoff, S.B., 2018. Differential Resting-State Connectivity Patterns of the Right Anterior and Posterior Dorsolateral Prefrontal Cortices (DLPFC) in Schizophrenia. Front. Psychiatry 9, 211. 10.3389/fpsyt.2018.00211

Cochran, S.M., Kennedy, M., McKerchar, C.E., Steward, L.J., Pratt, J.A., Morris, B.J., 2003. Induction of Metabolic Hypofunction and Neurochemical Deficits after Chronic Intermittent Exposure to Phencyclidine: Differential Modulation by Antipsychotic Drugs. Neuropsychopharmacology 28, 265–275. 10.1038/sj.npp.1300031

Condé, F., Maire□lepoivre, E., Audinat, E., Crépel, F., 1995. Afferent connections of the medial frontal cortex of the rat. II. Cortical and subcortical afferents. J. Comp. Neurol. 352, 567–593. 10.1002/cne.903520407

Coyle, J.T., 2012. NMDA Receptor and Schizophrenia: A Brief History. Schizophr. Bull. 38, 920–926. 10.1093/schbul/sbs076

d’Isa, R., Comi, G., Leocani, L., 2021. Apparatus design and behavioural testing protocol for the evaluation of spatial working memory in mice through the spontaneous alternation T-maze. Sci. Rep. 11, 21177. 10.1038/s41598-021-00402-7

Dandash, O., Fornito, A., Lee, J., Keefe, R.S.E., Chee, M.W.L., Adcock, R.A., Pantelis, C., Wood, S.J., Harrison, B.J., 2014. Altered Striatal Functional Connectivity in Subjects With an At-Risk Mental State for Psychosis. Schizophr. Bull. 40, 904–913. 10.1093/schbul/sbt093

Dawson, N., Kurihara, M., Thomson, D.M., Winchester, C.L., McVie, A., Hedde, J.R., Randall, A.D., Shen, S., Seymour, P.A., Hughes, Z.A., Dunlop, J., Brown, J.T., Brandon, N.J., Morris, B.J., Pratt, J.A., 2015. Altered functional brain network connectivity and glutamate system function in transgenic mice expressing truncated Disrupted-in-Schizophrenia 1. Transl. Psychiatry 5, e569–e569. 10.1038/tp.2015.60

Deiana, S., Hauber, W., Munster, A., Sommer, S., Ferger, B., Marti, A., Schmid, B., Dorner-Ciossek, C., Rosenbrock, H., 2022. Pro-cognitive effects of the GlyT1 inhibitor Bitopertin in rodents. Eur. J. Pharmacol. 935, 175306. 10.1016/j.ejphar.2022.175306

Driesen, N.R., McCarthy, G., Bhagwagar, Z., Bloch, M.H., Calhoun, V.D., D’Souza, D.C., Gueorguieva, R., He, G., Leung, H.-C., Ramani, R., Anticevic, A., Suckow, R.F., Morgan, P.T., Krystal, J.H., 2013. The Impact of NMDA Receptor Blockade on Human Working Memory-Related Prefrontal Function and Connectivity. Neuropsychopharmacology 38, 2613–2622. 10.1038/npp.2013.170

Ellison-Wright, I., Bullmore, E., 2009. Meta-analysis of diffusion tensor imaging studies in schizophrenia. Schizophr. Res. 108, 3–10. 10.1016/j.schres.2008.11.021

Fornito, A., Harrison, B.J., Goodby, E., Dean, A., Ooi, C., Nathan, P.J., Lennox, B.R., Jones, P.B., Suckling, J., Bullmore, E.T., 2013. Functional Dysconnectivity of Corticostriatal Circuitry as a Risk Phenotype for Psychosis. JAMA Psychiatry 70, 1143–1151. 10.1001/jamapsychiatry.2013.1976

Frohlich, J., Horn, J.D.V., 2014. Reviewing the ketamine model for schizophrenia. J Psychopharmacol 28, 287–302. 10.1177/0269881113512909

Fung, S.J., Webster, M.J., Sivagnanasundaram, S., Duncan, C., Elashoff, M., Weickert, C.S., 2010. Expression of Interneuron Markers in the Dorsolateral Prefrontal Cortex of the Developing Human and in Schizophrenia. Am. J. Psychiatry 167, 1479–1488. 10.1176/appi.ajp.2010.09060784

Gerhard, D.M., Pothula, S., Liu, R.-J., Wu, M., Li, X.-Y., Girgenti, M.J., Taylor, S.R., Duman, C.H., Delpire, E., Picciotto, M., Wohleb, E.S., Duman, R.S., 2019. GABA interneurons are the cellular trigger for ketamine’s rapid antidepressant actions. J. Clin. Investig. 130, 1336–1349. 10.1172/jci130808

Gonzalez-Burgos, G., Hashimoto, T., Lewis, D.A., 2010. Alterations of Cortical GABA Neurons and Network Oscillations in Schizophrenia. Curr. Psychiatry Rep. 12, 335–344. 10.1007/s11920-010-0124-8

Gonzalez-Burgos, G., Lewis, D.A., 2012. NMDA Receptor Hypofunction, Parvalbumin-Positive Neurons, and Cortical Gamma Oscillations in Schizophrenia. Schizophr. Bull. 38, 950–957. 10.1093/schbul/sbs010

Gur, R.E., Mozley, P.D., Resnick, S.M., Mozley, L.H., Shtasel, D.L., Gallacher, F., Arnold, S.E., Karp, J.S., Alavi, A., Reivich, M., Gur, R.C., 1995. Resting Cerebral Glucose Metabolism in First-Episode and Previously Treated Patients With Schizophrenia Relates to Clinical Features. Arch. Gen. Psychiatry 52, 657–667. 10.1001/archpsyc.1995.03950200047013

Hamm, J.P., Gilmore, C.S., Clementz, B.A., 2012. Augmented gamma band auditory steady-state responses: Support for NMDA hypofunction in schizophrenia. Schizophr. Res. 138, 1–7. 10.1016/j.schres.2012.04.003

Hashimoto, T., Volk, D.W., Eggan, S.M., Mirnics, K., Pierri, J.N., Sun, Z., Sampson, A.R., Lewis, D.A., 2003. Gene Expression Deficits in a Subclass of GABA Neurons in the Prefrontal Cortex of Subjects with Schizophrenia. J. Neurosci. 23, 6315–6326. 10.1523/jneurosci.23-15-06315.2003

Hill, K., Mann, L., Laws, K.R., Stephenson, C.M.E., Nimmo□Smith, I., McKenna, P.J., 2004. Hypofrontality in schizophrenia: a meta □ analysis of functional imaging studies. Acta Psychiatr. Scand. 110, 243–256. 10.1111/j.1600-0447.2004.00376.x

Homayoun, H., Moghaddam, B., 2007. NMDA Receptor Hypofunction Produces Opposite Effects on Prefrontal Cortex Interneurons and Pyramidal Neurons. J Neurosci 27, 11496– 11500. 10.1523/jneurosci.2213-07.2007

Howes, O.D., Kapur, S., 2009. The Dopamine Hypothesis of Schizophrenia: Version III— The Final Common Pathway. Schizophr. Bull. 35, 549–562. 10.1093/schbul/sbp006

Hwang, E., Brown, R.E., Kocsis, B., Kim, T., McKenna, J.T., McNally, J.M., Han, H.-B., Choi, J.H., 2019. Optogenetic stimulation of basal forebrain parvalbumin neurons modulates the cortical topography of auditory steady-state responses. Brain Struct. Funct. 224, 1505–1518. 10.1007/s00429-019-01845-5

Ionescu, T.M., Grohs-Metz, G., Hengerer, B., 2023. Functional ultrasound detects frequency-specific acute and delayed S-ketamine effects in the healthy mouse brain. Front. Neurosci. 17, 1177428. 10.3389/fnins.2023.1177428

Javitt, D.C., Zukin, S.R., Heresco-Levy, U., Umbricht, D., 2012. Has an Angel Shown the Way? Etiological and Therapeutic Implications of the PCP/NMDA Model of Schizophrenia. Schizophr. Bull. 38, 958–966. 10.1093/schbul/sbs069

Jodo, E., 2013. The role of the hippocampo-prefrontal cortex system in phencyclidine-induced psychosis: A model for schizophrenia. J. Physiol.-Paris 107, 434–440. 10.1016/j.jphysparis.2013.06.002

Jonckers, E., Palacios, R.D. y, Shah, D., Guglielmetti, C., Verhoye, M., Linden, A., 2014. Different anesthesia regimes modulate the functional connectivity outcome in mice. Magn. Reson. Med. 72, 1103–1112. 10.1002/mrm.24990

Jonckers, E., Shah, D., Hamaide, J., Verhoye, M., Linden, A.V. der, 2015. The power of using functional fMRI on small rodents to study brain pharmacology and disease. Front Pharmacol 6, 231. 10.3389/fphar.2015.00231

Karcher, N.R., Rogers, B.P., Woodward, N.D., 2019. Functional Connectivity of the Striatum in Schizophrenia and Psychotic Bipolar Disorder. Biol. Psychiatry: Cogn. Neurosci. Neuroimaging 4, 956–965. 10.1016/j.bpsc.2019.05.017

Keilhoff, G., Becker, A., Grecksch, G., Wolf, G., Bernstein, H.-G., 2004. Repeated application of ketamine to rats induces changes in the hippocampal expression of parvalbumin, neuronal nitric oxide synthase and cFOS similar to those found in human schizophrenia. Neuroscience 126, 591–598. 10.1016/j.neuroscience.2004.03.039

Kim, S., Jang, S.-K., Kim, D.-W., Shim, M., Kim, Y.-W., Im, C.-H., Lee, S.-H., 2019. Cortical volume and 40-Hz auditory-steady-state responses in patients with schizophrenia and healthy controls. NeuroImage: Clin. 22, 101732. 10.1016/j.nicl.2019.101732

Kokkinou, M., Irvine, E.E., Bonsall, D.R., Natesan, S., Wells, L.A., Smith, M., Glegola, J., Paul, E.J., Tossell, K., Veronese, M., Khadayate, S., Dedic, N., Hopkins, S.C., Ungless, M.A., Withers, D.J., Howes, O.D., 2021. Reproducing the dopamine pathophysiology of schizophrenia and approaches to ameliorate it: a translational imaging study with ketamine. Mol. Psychiatry 26, 2562–2576. 10.1038/s41380-020-0740-6

Kotoula, V., Webster, T., Stone, J., Mehta, M.A., 2021. Resting-state connectivity studies as a marker of the acute and delayed effects of subanaesthetic ketamine administration in healthy and depressed individuals: A systematic review. Brain Neurosci Adv 5, 23982128211055424. 10.1177/23982128211055426

Krystal, J.H., Karper, L.P., Seibyl, J.P., Freeman, G.K., Delaney, R., Bremner, J.D., Heninger, G.R., Bowers, M.B., Charney, D.S., 1994. Subanesthetic Effects of the Noncompetitive NMDA Antagonist, Ketamine, in Humans: Psychotomimetic, Perceptual, Cognitive, and Neuroendocrine Responses. Arch. Gen. Psychiatry 51, 199–214. 10.1001/archpsyc.1994.03950030035004

Krzystanek, M., Pałasz, A., 2019. NMDA Receptor Model of Antipsychotic Drug-Induced Hypofrontality. Int. J. Mol. Sci. 20, 1442. 10.3390/ijms20061442

Lee, G., Zhou, Y., 2019. NMDAR Hypofunction Animal Models of Schizophrenia. Front Mol Neurosci 12, 185. 10.3389/fnmol.2019.00185

Leishman, E., O’Donnell, B.F., Millward, J.B., Vohs, J.L., Rass, O., Krishnan, G.P., Bolbecker, A.R., Morzorati, S.L., 2015. Phencyclidine Disrupts the Auditory Steady State Response in Rats. PLoS ONE 10, e0134979. 10.1371/journal.pone.0134979

Lewis, D.A., Curley, A.A., Glausier, J.R., Volk, D.W., 2012. Cortical parvalbumin interneurons and cognitive dysfunction in schizophrenia. Trends Neurosci. 35, 57–67. 10.1016/j.tins.2011.10.004

Lewis, D.A., Hashimoto, T., Volk, D.W., 2005. Cortical inhibitory neurons and schizophrenia. Nat. Rev. Neurosci. 6, 312–324. 10.1038/nrn1648

Li, N., Lee, B., Liu, R.-J., Banasr, M., Dwyer, J.M., Iwata, M., Li, X.-Y., Aghajanian, G., Duman, R.S., 2010. mTOR-Dependent Synapse Formation Underlies the Rapid Antidepressant Effects of NMDA Antagonists. Science 329, 959–964. 10.1126/science.1190287

Light, G.A., Hsu, J.L., Hsieh, M.H., Meyer-Gomes, K., Sprock, J., Swerdlow, N.R., Braff, D.L., 2006. Gamma Band Oscillations Reveal Neural Network Cortical Coherence Dysfunction in Schizophrenia Patients. Biol. Psychiatry 60, 1231–1240. 10.1016/j.biopsych.2006.03.055

Lisman, J.E., Fellous, J.-M., Wang, X.-J., 1998. A role for NMDA-receptor channels in working memory. Nat. Neurosci. 1, 273–275. 10.1038/1086

Manoach, D.S., Gollub, R.L., Benson, E.S., Searl, M.M., Goff, D.C., Halpern, E., Saper, C.B., Rauch, S.L., 2000. Schizophrenic subjects show aberrant fMRI activation of dorsolateral prefrontal cortex and basal ganglia during working memory performance. Biol. Psychiatry 48, 99–109. 10.1016/s0006-3223(00)00227-4

Manoach, D.S., Press, D.Z., Thangaraj, V., Searl, M.M., Goff, D.C., Halpern, E., Saper, C.B., Warach, S., 1999. Schizophrenic subjects activate dorsolateral prefrontal cortex during a working memory task, as measured by fMRI. Biol. Psychiatry 45, 1128–1137. 10.1016/s0006-3223(98)00318-7

Marek, R., Jin, J., Goode, T.D., Giustino, T.F., Wang, Q., Acca, G.M., Holehonnur, R., Ploski, J.E., Fitzgerald, P.J., Lynagh, T., Lynch, J.W., Maren, S., Sah, P., 2018. Hippocampus-driven feed-forward inhibition of the prefrontal cortex mediates relapse of extinguished fear. Nat Neurosci 21, 384–392. 10.1038/s41593-018-0073-9

Marín, O., 2012. Interneuron dysfunction in psychiatric disorders. Nat. Rev. Neurosci. 13, 107–120. 10.1038/nrn3155

Masaki, Y., Kashiwagi, Y., Watabe, H., Abe, K., 2019. (R) □ and (S) □ ketamine induce differential fMRI responses in conscious rats. Synapse 73, e22126. 10.1002/syn.22126

Mellios, N., Huang, H.-S., Baker, S.P., Galdzicka, M., Ginns, E., Akbarian, S., 2009. Molecular Determinants of Dysregulated GABAergic Gene Expression in the Prefrontal Cortex of Subjects with Schizophrenia. Biol. Psychiatry 65, 1006–1014. 10.1016/j.biopsych.2008.11.019

Meyer-Lindenberg, A., Miletich, R.S., Kohn, P.D., Esposito, G., Carson, R.E., Quarantelli, M., Weinberger, D.R., Berman, K.F., 2002. Reduced prefrontal activity predicts exaggerated striatal dopaminergic function in schizophrenia. Nat. Neurosci. 5, 267–271. 10.1038/nn804

Meyer-Lindenberg, A., Weinberger, D.R., 2006. Intermediate phenotypes and genetic mechanisms of psychiatric disorders. Nat. Rev. Neurosci. 7, 818–827. 10.1038/nrn1993

Moghaddam, B., 2003. Bringing Order to the Glutamate Chaos in Schizophrenia. Neuron 40, 881–884. 10.1016/s0896-6273(03)00757-8

Moghaddam, B., Adams, B., Verma, A., Daly, D., 1997. Activation of Glutamatergic Neurotransmission by Ketamine: A Novel Step in the Pathway from NMDA Receptor Blockade to Dopaminergic and Cognitive Disruptions Associated with the Prefrontal Cortex. J. Neurosci. 17, 2921–2927. 10.1523/jneurosci.17-08-02921.1997

Moghaddam, B., Adams, B.W., 1998. Reversal of Phencyclidine Effects by a Group II Metabotropic Glutamate Receptor Agonist in Rats. Science 281, 1349–1352. 10.1126/science.281.5381.1349

Montani, C., Canella, C., Schwarz, A.J., Li, J., Gilmour, G., Galbusera, A., Wafford, K., Gutierrez-Barragan, D., McCarthy, A., Shaw, D., Knitowski, K., McKinzie, D., Gozzi, A., Felder, C., 2021. The M1/M4 preferring muscarinic agonist xanomeline modulates functional connectivity and NMDAR antagonist-induced changes in the mouse brain. Neuropsychopharmacology 46, 1194–1206. 10.1038/s41386-020-00916-0

Mukai, J., Tamura, M., Fénelon, K., Rosen, A.M., Spellman, T.J., Kang, R., MacDermott, A.B., Karayiorgou, M., Gordon, J.A., Gogos, J.A., 2015. Molecular Substrates of Altered Axonal Growth and Brain Connectivity in a Mouse Model of Schizophrenia. Neuron 86, 680–695. 10.1016/j.neuron.2015.04.003

Murray, A.J., Woloszynowska-Fraser, M.U., Ansel-Bollepalli, L., Cole, K.L.H., Foggetti, A., Crouch, B., Riedel, G., Wulff, P., 2015. Parvalbumin-positive interneurons of the prefrontal cortex support working memory and cognitive flexibility. Sci. Rep. 5, 16778. 10.1038/srep16778

Nakazawa, K., Jeevakumar, V., Nakao, K., 2017. Spatial and temporal boundaries of NMDA receptor hypofunction leading to schizophrenia. npj Schizophr. 3, 7. 10.1038/s41537-016-0003-3

Orduz, D., Bischop, D.P., Schwaller, B., Schiffmann, S.N., Gall, D., 2013. Parvalbumin tunes spike □ timing and efferent short □ term plasticity in striatal fast spiking interneurons. J. Physiol. 591, 3215–3232. 10.1113/jphysiol.2012.250795

Orliac, F., Naveau, M., Joliot, M., Delcroix, N., Razafimandimby, A., Brazo, P., Dollfus, S., Delamillieure, P., 2013. Links among resting-state default-mode network, salience network, and symptomatology in schizophrenia. Schizophr. Res. 148, 74–80. 10.1016/j.schres.2013.05.007

Parent, M.A., Wang, L., Su, J., Netoff, T., Yuan, L.-L., 2010. Identification of the Hippocampal Input to Medial Prefrontal Cortex In Vitro. Cereb. Cortex 20, 393–403. 10.1093/cercor/bhp108

Petreanu, L., Huber, D., Sobczyk, A., Svoboda, K., 2007. Channelrhodopsin-2–assisted circuit mapping of long-range callosal projections. Nat. Neurosci. 10, 663–668. 10.1038/nn1891

Powell, S.B., Weber, M., Geyer, M.A., 2012. Genetic Models of Sensorimotor Gating: Relevance to Neuropsychiatric Disorders. Curr. Top. Behav. Neurosci. 12, 251–318. 10.1007/7854_2011_195

Reilly, T.J., Nottage, J.F., Studerus, E., Rutigliano, G., Micheli, A.I.D., Fusar-Poli, P., McGuire, P., 2018. Gamma band oscillations in the early phase of psychosis: A systematic review. Neurosci. Biobehav. Rev. 90, 381–399. 10.1016/j.neubiorev.2018.04.006

Rezvani, A., 2006. Animal Models of Cognitive Impairment. Front. Neurosci. 20064835, 37– 48. 10.1201/9781420004335.ch4

Roach, B.J., Ford, J.M., Mathalon, D.H., 2019. Gamma Band Phase Delay in Schizophrenia. Biol. Psychiatry: Cogn. Neurosci. Neuroimaging 4, 131–139. 10.1016/j.bpsc.2018.08.011

Romón, T., Mengod, G., Adell, A., 2011. Expression of parvalbumin and glutamic acid decarboxylase-67 after acute administration of MK-801. Implications for the NMDA hypofunction model of schizophrenia. Psychopharmacology 217, 231–238. 10.1007/s00213-011-2268-6

Sarpal, D.K., Robinson, D.G., Lencz, T., Argyelan, M., Ikuta, T., Karlsgodt, K., Gallego, J.A., Kane, J.M., Szeszko, P.R., Malhotra, A.K., 2015. Antipsychotic Treatment and Functional Connectivity of the Striatum in First-Episode Schizophrenia. JAMA Psychiatry 72, 5–13. 10.1001/jamapsychiatry.2014.1734

Schuelert, N., Dorner□Ciossek, C., Brendel, M., Rosenbrock, H., 2018. A comprehensive analysis of auditory event □ related potentials and network oscillations in an NMDA receptor antagonist mouse model using a novel wireless recording technology. Physiological Reports 6, e13782. 10.14814/phy2.13782

Sigurdsson, T., 2016. Neural circuit dysfunction in schizophrenia: Insights from animal models. Neuroscience 321, 42–65. 10.1016/j.neuroscience.2015.06.059

Sigurdsson, T., Duvarci, S., 2016. Hippocampal-Prefrontal Interactions in Cognition, Behavior and Psychiatric Disease. Frontiers Syst Neurosci 9, 190. 10.3389/fnsys.2015.00190

Sivarao, D.V., Chen, P., Senapati, A., Yang, Y., Fernandes, A., Benitex, Y., Whiterock, V., Li, Y.-W., Ahlijanian, M.K., 2016. 40 Hz Auditory Steady-State Response Is a Pharmacodynamic Biomarker for Cortical NMDA Receptors. Neuropsychopharmacology 41, 2232–2240. 10.1038/npp.2016.17

Soyka, M., Koch, W., Möller, H.J., Rüther, T., Tatsch, K., 2005. Hypermetabolic pattern in frontal cortex and other brain regions in unmedicated schizophrenia patients. Eur. Arch. Psychiatry Clin. Neurosci. 255, 308–312. 10.1007/s00406-005-0563-0

Stone, J.M., Day, F., Tsagaraki, H., Valli, I., McLean, M.A., Lythgoe, D.J., O’Gorman, R.L., Barker, G.J., McGuire, P.K., OASIS, 2009. Glutamate Dysfunction in People with Prodromal Symptoms of Psychosis: Relationship to Gray Matter Volume. Biol. Psychiatry 66, 533–539. 10.1016/j.biopsych.2009.05.006

Strobel, B., Miller, F.D., Rist, W., Lamla, T., 2015. Comparative Analysis of Cesium Chloride- and Iodixanol-Based Purification of Recombinant Adeno-Associated Viral Vectors for Preclinical Applications. Hum. Gene Ther. Methods 26, 147–157. 10.1089/hgtb.2015.051

Strobel, B., Zuckschwerdt, K., Zimmermann, G., Mayer, C., Eytner, R., Rechtsteiner, P., Kreuz, S., Lamla, T., 2019. Standardized, Scalable, and Timely Flexible Adeno-Associated Virus Vector Production Using Frozen High-Density HEK-293 Cell Stocks and CELLdiscs. Hum. Gene Ther. Methods 30, 23–33. 10.1089/hgtb.2018.228

Suzuki, Y., Jodo, E., Takeuchi, S., Niwa, S., Kayama, Y., 2002. Acute administration of phencyclidine induces tonic activation of medial prefrontal cortex neurons in freely moving rats. Neuroscience 114, 769–779. 10.1016/s0306-4522(02)00298-1

Tannenbaum, J., Bennett, B.T., 2015. Russell and Burch’s 3Rs then and now: the need for clarity in definition and purpose. J. Am. Assoc. Lab. Anim. Sci. □: JAALAS 54, 120–32.

Theberge, J., Bartha, R., Drost, D.J., Menon, R.S., Malla, A., Takhar, J., Neufeld, R.W., Rogers, J., Pavlosky, W., Schaefer, B., Densmore, M., Al-Semaan, Y., Williamson, P.C., 2002. Glutamate and Glutamine Measured With 4.0 T Proton MRS in Never-Treated Patients With Schizophrenia and Healthy Volunteers. Am. J. Psychiatry 159, 1944–1946. 10.1176/appi.ajp.159.11.1944

Toader, O., Heimendahl, M. von, Schuelert, N., Nissen, W., Rosenbrock, H., 2020. Suppression of Parvalbumin Interneuron Activity in the Prefrontal Cortex Recapitulates Features of Impaired Excitatory/Inhibitory Balance and Sensory Processing in Schizophrenia. Schizophr. Bull. 46, 981–989. 10.1093/schbul/sbz123

Tost, H., Bilek, E., Meyer-Lindenberg, A., 2012. Brain connectivity in psychiatric imaging genetics. NeuroImage 62, 2250–2260. 10.1016/j.neuroimage.2011.11.007

Townsend, L., Pillinger, T., Selvaggi, P., Veronese, M., Turkheimer, F., Howes, O., 2022. Brain glucose metabolism in schizophrenia: a systematic review and meta-analysis of 18 FDG-PET studies in schizophrenia. Psychol. Med. 1–18. 10.1017/s003329172200174x

Zhang, Z.J., Reynolds, G.P., 2002. A selective decrease in the relative density of parvalbumin-immunoreactive neurons in the hippocampus in schizophrenia. Schizophr. Res. 55, 1–10. 10.1016/s0920-9964(01)00188-8

Zhou, Z., Zhang, G., Li, X., Liu, X., Wang, N., Qiu, L., Liu, W., Zuo, Z., Yang, J., 2015. Loss of Phenotype of Parvalbumin Interneurons in Rat Prefrontal Cortex Is Involved in Antidepressant- and Propsychotic-Like Behaviors Following Acute and Repeated Ketamine Administration. Mol. Neurobiol. 51, 808–819. 10.1007/s12035-014-8798-2

