## Supplemental Info for "Long-term adaptation of prefrontal circuits in a mouse model of NMDAR hypofunction"

### SUPPLEMENTARY MATERIALS AND METHODS

#### Functional ultrasound imaging

At the beginning of each scan, the animals were sedated using 4% isoflurane in oxygen. Afterwards, the concentration of isoflurane was decreased and maintained at 1% before the animals were positioned in a stereotactic frame (David Kopf Instruments, USA). A catheter was inserted subcutaneously for dexmedetomidine constant infusion. During all sessions, the temperature was maintained at 37.0° C (PhysioSuite homeothermic warming system, Kent Scientific Corporation, USA). Additionally, heart rates and blood oxygenation levels were constantly monitored using a pulse oximeter placed on the hind paw. Prior to the scans, the animals were injected subcutaneously with 0.05 mg/kg meloxicam (Metacam®, Boehringer Ingelheim, DE) to minimize discomfort. Sedation was switched to dexmedetomidine by first applying a bolus of 0.1 mg/kg dexmedetomidine hydrochloride (Tocris, USA) followed by a constant subcutaneous dexmedetomidine infusion (0.2 mg/kg/h, 5 mL/kg/h flow) throughout the session, while simultaneously decreasing the delivered isoflurane by 0.1% per minute until complete discontinuation. The scalps of the mice were shaved before being covered by ultrasonic gel. Next, we lowered the ultrasonic probe to 1mm above the scalp, being completely immersed in the ultrasound gel. The probe was composed of 128 piezoelectric transducers (0.08mm per element), emitting and receiving backscattered echoes at a central frequency of 15MHz. For imaging, we used a 128-channel scanner (Iconeus One, Iconeus, Paris, France), generating images using 200 compounded frames acquired at a frame rate of 500Hz. Each final image generated by summing echoes at 11 different angles -10° and 10° with separations of 2°[45–49]. First, an angiographic image was acquired by imaging 30 slices with 0.2mm between-slice separation, covering the brain between midbrain and frontal cortex. Functional ultrasound was performed in a single left-hemispheric oblique slice, identical to the one used in our previous work[47] and comprising various cortical and subcortical images listed in Fig. 5. To this extent, the Brain Positioning System[50] (Iconeus, France) was used to correctly position the probe via the angiographic image acquired for the respective animal. Functional ultrasound imaging was not initiated earlier than 10 minutes after we discontinued the isoflurane administration, the measurements lasting for 60 minutes, thereby comprising 9000 2D frames at a resolution of 2.5Hz. To isolate blood signal from the images, a spatiotemporal singular value decomposition clutter filter was employed[45]. Please refer to[51] for a detailed report of how the images were generated. After each session, to antagonize the dexmedetomidine effects, we injected the mice using a five times higher dose of atipamezole (Alzane, Zoetis, Germany) compared to the total applied dexmedetomidine.

For data analysis, Regional Power Doppler time courses were extracted from 12 different brain areas in the left hemisphere. The raw Power Doppler signals were then scrubbed of artifacts using a protocol adapted from[52]. Next, the data were temporally filtered using a second-order Butterworth filter to a bandpass between 0.01 and 0.2Hz.

### Supplementary Figures and Tables

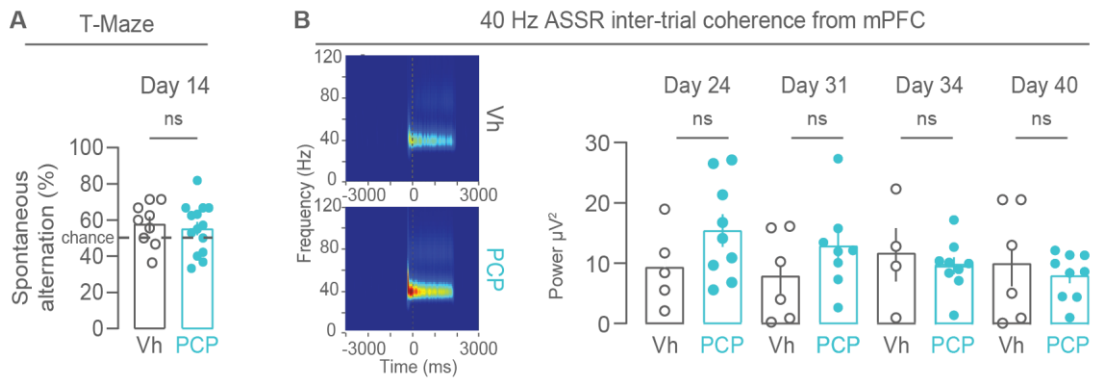


**Sup. Fig. 1:** A Percent of spontaneous alternation of vehicle- and PCP treated mice in the T-Maze task at day 14 (for Vh, mean spontaneous alternation: 55.9 ± 4 %, n= 9 mice; for PCP: 55.4 ± 3.7%, n= 14 mice; Mann-Whitney U test: U = 55.5, p = 0.65)**. B** Heatmaps of power of 40 Hz ASSR from the mPFC of vehicle- and PCP-treated animals at Day 24 (left) and quantifications (right) at day 24 (for Vh, mean power: 9.22 ± 2.9, n= 5 mice; for PCP: 15.3 ± 2.7, n= 9 mice; Mann-Whitney U test: U = 12, p = 0.189); day 31 (for Vh, mean power: 7.78 ± 2.95, n= 6 mice; for PCP: 12.77 ± 2.54, n= 8 mice; Mann-Whitney U test: U = 16, p = 0.345); day 34 (for Vh, mean power: 11.57± 4.41, n= 4 mice; for PCP: 9.74 ± 1.41, n= 9 mice; Mann-Whitney U test: U = 15, p = 0.355) and day 40 (for Vh, mean power: 10.18 ± 3.81, n= 6 mice; for PCP: 8.139 ± 1.28, n= 9 mice; Mann-Whitney U test: U = 24, p = 0.775). Each dot is the quantification of a single animal.

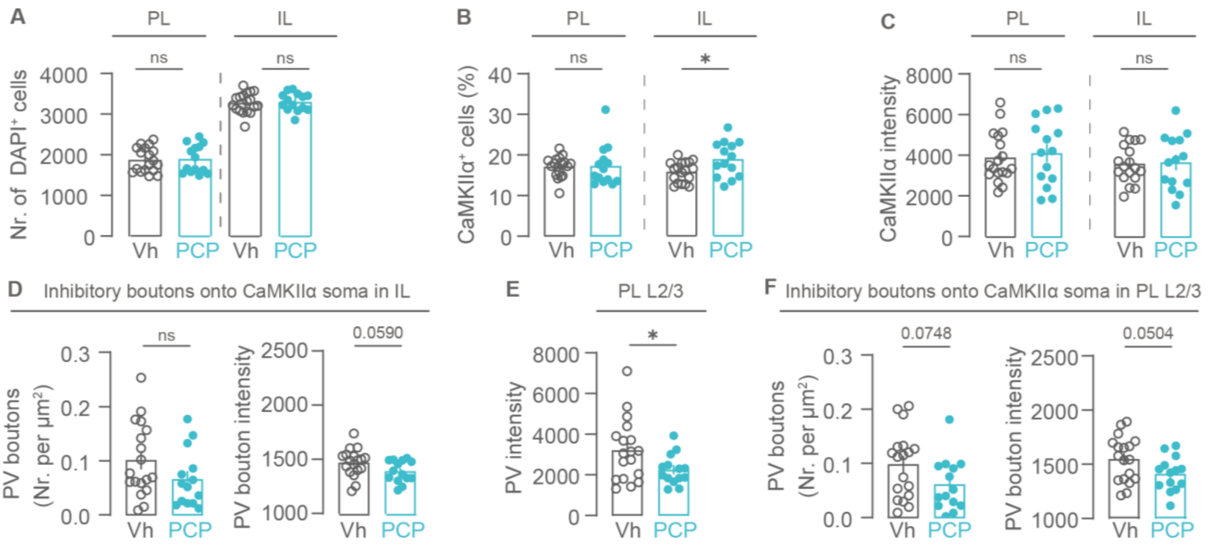


**Suppl. Fig.2: A** Total number of DAPI$^+$ cell nuclei detected in the PL and IL of vehicle- and PCP-treated mice (PL: for Vh, mean total count: 1874 ± 70.94, n=18 sections from 9 mice; For PCP, mean total count: 1897 ± 93.29, n=14 sections from 7 mice; 2-tailed unpaired Mann-Whitney U-test, U = 121, p = 0.8587; IL: for Vh, mean total count: 3275 ± 59.09, n=18 sections from 9 mice; For PCP, mean total count: 3298 ± 59.37, n=14 sections from 7 mice; 2-tailed unpaired t-test: t(30) = 0.2646, p = 0.7932). **B** Percental count of CaMKIIα cells in PL (left) and IL (right) (PL: for Vh, mean total count: 17.08 ± 0.60, n=18 sections from 9 mice; For PCP, mean total count: 17.25 ± 1.33, n=14 sections from 7 mice; 2-tailed Mann-Whitney U-test: U = 104, p = 0.4195; IL: for Vh, mean total count: 15.56 ± 0.59, n=18 sections from 9 mice; For PCP, mean total count: 18.67 ± 1.16, n=14 sections from 7 mice; 2-tailed unpaired t-test: t(30) = 2.545, p = 0.0163). **C** Fluorescent intensity of CaMKIIα$^+$ cells in PL (left) and IL (right) (PL: for Vh, mean intensity: 3885 ± 292.8, n=18 sections from 9 mice; For PCP, mean intensity: 4097 ± 423.3, n=14 sections from 7 mice; 2-tailed unpaired t-test: t(30) = 0.4247, p = 0.6741; IL: for Vh, mean intensity: 3609 ± 220.5, n=18 sections from 9 mice; For PCP, mean intensity: 3675 ± 361.5, n=14 sections from 7 mice; 2-tailed unpaired t-test: t(30) = 0.1622, p = 0.8723). **D** Left: Density of inhibitory PV$^+$ boutons on the soma of CaMKIIα$^+$ excitatory cells in IL of Vh- and PCP-treated mice (for Vh, mean density: 0.1008 ± 0.02, n=18 sections from 9 mice; For PCP, mean density: 0.06602 ± 0.01, n=14 sections from 7 mice; 2-tailed unpaired Mann-Whitney U-test, U = 86, p = 0.1350). Right: PV fluorescence intensity of PV$^+$ boutons on CaMKIIα somata in IL (for Vh, mean intensity: 1469 ± 29.78, n=18 sections from 9 mice; For PCP, mean intensity: 1388 ± 26.68, n=14 sections from 7 mice; 2-tailed unpaired t-test: t(30) = 1.963, p = 0.0590). **E** Fluorescence intensity of PV$^+$ cells in Layer 2/3 of PL (for Vh, mean intensity: 3208 ± 361.9, n=18 sections from 9 mice; For PCP, mean intensity: 2245 ± 198.4, n=14 sections from 7 mice; 2-tailed unpaired t-test: t (30) = 2.154, p = 0.0394). **F** Left: Density of inhibitory PV$^+$ boutons on the soma of CaMKIIα$^+$ excitatory cells in Layer 2/3 of PL (for Vh, mean density: 0.09839 ± 0.01, n=18 sections from 9 mice; For PCP, mean density: 0.06106 ± 0.01, n=14 sections from 7 mice; 2-tailed unpaired t-test: t(30) = 1.846, p = 0.0748). Right: PV fluorescence intensity of Layer 2/3 PV$^+$ boutons on CaMKIIα somata in IL (for Vh, mean intensity: 1547 ± 49.35, n=18 sections from 9 mice; For PCP, mean intensity: 1410 ± 42.03, n=14 sections from 7 mice; 2-tailed unpaired t-test: t(30) = 2.038, p = 0.0504). Each data point represents mean measured per brain slice.

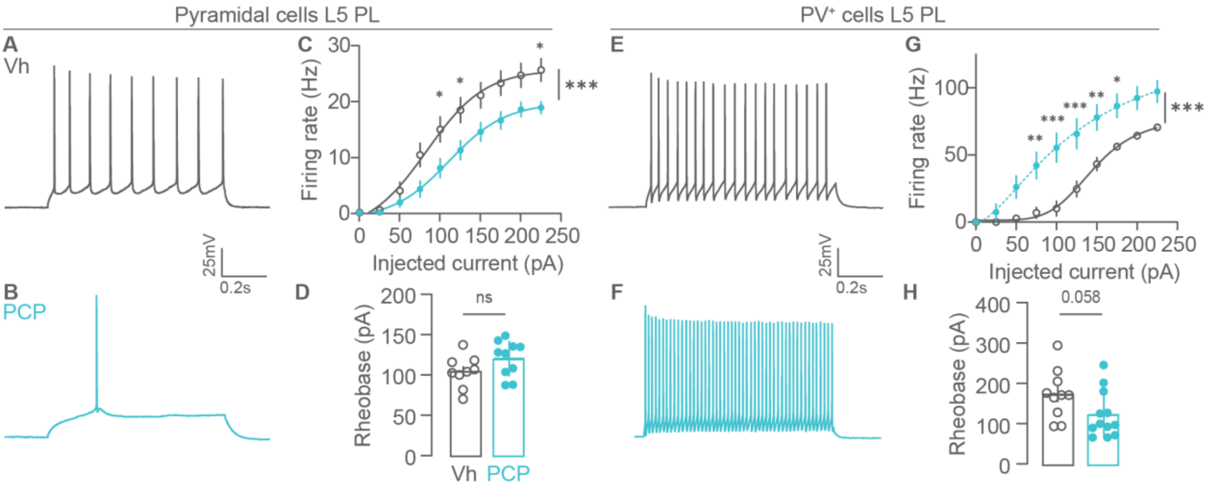


**Suppl. Fig. 3: A-B** Representative traces of spikes evoked by a 75 pA depolarizing current pulse in pyramidal cells of the L5 PL in vehicle- (**A**) and PCP-treated mice (**B**). **C** Frequency versus injected current amplitude for recorded pyramidal cells from vehicle- and PCP treated animals (two-way repeated measures ANOVA, Current injected x Treatment interaction: F (9, 153) = 4.771, P<0.0001 followed by Šídák's multiple comparisons test; for Vh: n= 8 cells from 6 mice and for PCP: n= 11 cells from 4 mice). **D** Rheobase for recorded pyramidal cells from vehicle- and PCP-treated animals (For Vh, mean rheobase: 103 ± 6.6pA, n= 9 cells from 7 mice; For PCP, mean rheobase: 119.3 ± 7pA, n= 10 cells from 5 mice; 2-tailed unpaired t test: t(17) = 1.68, p = 0.111). **E-F** Representative traces of spikes evoked by a 75 pA depolarizing current pulse in PV$^+$ interneurons of the L5 PL in vehicle- (**E**) and PCP-treated mice (**F**) Intrinsic properties of PV+ interneurons were assessed using whole-cell patch clamp recording from mCherry-expressing cells in slices of PV-tdTomato mice. **G** Frequency versus injected current amplitude for recorded PV$^+$ interneurons from vehicle- and PCP treated animals (two-way repeated measures ANOVA, Current injected x Treatment interaction: F (9, 171) = 7.909, P<0.0001 followed by Šídák's multiple comparisons test; for Vh: n= 10 cells from 4 mice and for PCP: n= 11 cells from 4 mice). **H** Rheobase for recorded PV$^+$ interneurons from vehicle- and PCP treated animals (For Vh, mean rheobase: 172.7 ± 19.3pA, n=11 cells from 4 mice; For PCP, mean rheobase: 124.5 ± 18 pA, n= 12 cells from 5 mice; 2-tailed unpaired t-test: t(20) = 2.01, p = 0.0581). Each dot is the quantification of a single cell.

| | **Vehicle**<br>n = 9 cells from 7 mice | **PCP**<br>n = 10 cells from 5 mice |
|---|---|---|
| RMP (mV) | -62.6 ± 2.1 | -62.2 ± 0.7 |
| $R_I$ (MΩ) | 153.2 ± 10.1 | 135.7 ± 11.1 |
| Sag ratio | 0.17 ± 0.04 | 0.22 ± 0.03 |
| <u>Action potential:</u> | | |
| Peak (mV) | 102.45 ± 2.2 | 102.4 ± 1.7 |
| Half-Width (ms) | 1.52 ± 0.06 | 1.45± 0.04 |
| Threshold (mV) | -42.7 ± 1.4 | -40.7 ± 1.0 |

**Suppl. Table 1: Passive and active membrane properties of L5 pyramidal cells in the mPFC of vehicle and PCP treated animals.** *RMP, resting membrane potential; $R_I$, input resistance. The properties of action potentials were recorded starting from the current injection step that triggered the first action potential. Properties were measured for the first action potential only. No significant difference between vehicle and PCP-treated animals were observed in any parameter (P>0.1). 2-tailed unpaired t-test for all except for Sag ratio: Mann-Whitney U test. Data shown as mean ± S.E.M.*

| | **Vehicle**<br>n = 11 cells from 4 mice | **PCP**<br>n = 12 cells from 5 mice |
|---|---|---|
| RMP (mV) | -60.2 ± 1.6 | -64.4 ± 1*<br>2-tailed unpaired t-test t(18) = 2.438, p= 0.0254 |
| $R_I$ (MΩ) | 159.2 ± 9.4 | 187.05 ± 18.3 |
| Sag ratio | 0.09 ± 0.01 | 0.1 ± 0.01 |
| <u>Action potential:</u> | | |
| Peak (mV) | 88.8 ± 2.1 | 94.3 ± 2.6 |
| Half-Width (ms) | 0.58 ± 0.02 | 0.53 ± 0.03 |
| Threshold (mV) | -43.4 ± 1.9 | -48.2 ± 2.3 |

**Suppl. Table 2: Passive and active membrane properties of L5 PL PV+ interneurons in the mPFC of vehicle and PCP treated animals.** *RMP, resting membrane potential; $R_I$, input resistance. The properties of action potentials were recorded starting from the current injection step that triggered the first action potential. Properties were measured for the first action potential only. No significant difference between vehicle and PCP-treated animals were observed in any parameter (P>0.1) except for the RMP. 2-tailed unpaired t-test for all except for Sag ratio: Mann-Whitney U test. Data shown as mean ± S.E.M.*

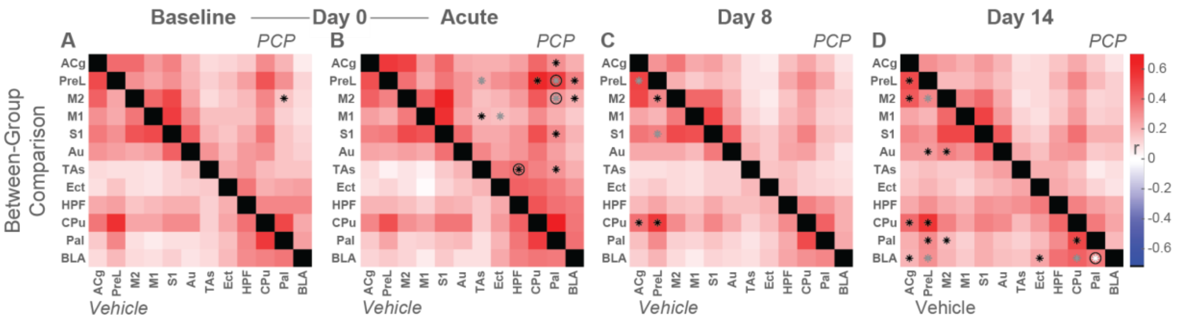


**Suppl. Fig. 4: A-D** Comparison of functional connectivity matrices of 12 regions covered by the acquired slice (below diagonal: vehicle cohort, above diagonal: PCP cohort) at **A** day 0 – baseline (minutes 5-15), **B** acute (minutes 25-35), **C** day 8 (minutes 5-60), and **D** day 14 (minutes 5-60). Matrices show the Z-transformed average Pearsons'r correlation coefficients for the two cohorts along with connections showing significant differences at the respective timepoints (*black: $p < 0.05$, *gray: $p < 0.01$, *white: $p < 0.001$, O: $p < 0.05$, FDR-corrected, two-sample t-tests).
